# Granulosa cell glycogen fuels the avascular corpus luteum

**DOI:** 10.1101/2025.01.22.634063

**Authors:** Jianning Liao, Qinghua Liu, Cong Liu, Guiqiong Liu, Xiang Li, Xiaodong Wang, Yaqin Wang, Ruiyan Liu, Hao Wu, Chaoli Wang, Hongru Shi, Yongheng Zhao, Wenkai Ke, Zaohong Ran, Zian Wu, Bowen Tan, Quanfeng Wang, Guohua Hua, Shujun Zhang, Qingzhen Xie, Guoshi Liu, Changjiu He

**Affiliations:** Key Laboratory of Agricultural Animal Genetics, Breeding and Reproduction of Ministry of Education, College of Animal Sciences and Technology, Huazhong Agricultural University, Wuhan 430070, PR China; Centre for Reproductive Medicine, Renmin Hospital of Wuhan University, Wuhan 430000, China; Key Laboratory of Animal Genetics and Breeding of the Ministry of Agriculture, College of Animal Science and Technology, China Agricultural University, Beijing, China; College of Animal Science and Technology, Shihezi University, Shihezi, China; Xinjiang Western Animal Husbandry Co., Ltd, Shihezi, China; Xinjiang Jinken aoqun agriculture and animal husbandry technology Co., Ltd, Hetian, China; Binzhou Bincheng District Animal Husbandry and Veterinary Service Center, Binzhou, China

**Keywords:** ovulation, glycogen, granulosa cell, corpus luteum, follicle, luteinization

## Abstract

The corpus luteum (CL) arises from the luteinization of follicular granulosa cells (GCs) and theca cells, marked by rapid progesterone elevation and angiogenesis. Intriguingly, angiogenesis lags behind progesterone elevation, creating an avascular phase during which luteal cells must fuel intensive steroidogenesis without perfusion. How the avascular CL meets this energetic demand remains a mystery. Here, we reveal a novel cellular adaptive mechanism–GC energy storage (GCES)–that resolves this enigma. We demonstrate that upon luteinization initiation, GCs enter a metabolically quiescent state yet enhance glucose uptake via SLC2A1, converting the glucose into glycogen through the hCG (LH)-MAPK-RUNX1-Insulin signaling axis. Catabolism of this glycogen reserve supplies the energy required for the avascular CL, ensuring normal luteogenesis. GCES is evolutionarily conserved across species. Genetic or pharmacological disruption of GCES or glycogenolysis induces luteal insufficiency, whereas timely glucose administration enhances GCES, improving luteal function and optimize reproductive outcome in both mouse and ovine models. In human study, orally intake of glucose post-hCG significantly augments GCES and enhances progesterone production in women. These results advance luteal physiology by uncovering a universal reproductive principle with direct clinical implications.

## INTRODUCTION

The corpus luteum (CL) is a temporary ovarian endocrine structure critical for reproduction. It sustains pregnancy by secreting progesterone and orchestrates the reproductive cycle via its cyclic formation and regression [1, 2]. Luteal insufficiency, defined by the suboptimal progesterone production and shortened luteal phase, represents a common etiology of female subfertility and is closely linked to reproductive disorders, including menstrual disorders, premenstrual spotting and recurrent pregnancy loss [3, 4]. It is estimated that ∼8% of reproductive-age women experience luteal insufficiency, with the incidences escalating in parallel to maternal age [3, 5–8]. Notably, luteal insufficiency is prevalent in recurrent miscarriage cases [9, 10], underscoring its profound impact on reproductive health.

CL arises from the luteinization of granulosa cells (GCs) and theca cells, a process triggered by the pre-ovulatory surge of luteinizing hormone (LH) or human chorionic gonadotropin (hCG, an LH-equivalent hormone) [11]. Luteinization encompasses multiple coordinated cellular mechanisms, such as proliferation cessation, steroidogenesis reprogramming, GC-layer stiffening, cell migration and neovascularization [2, 12–15]. LH (hCG)-activated downstream regulators, including P21CIP1, P27KIP1, ERK1/2, PI3K/AKT, CEBPα/β, RUNX1/2, NR4A2/NURR1, CREB, cortisol/glucocorticoid receptors and H3K27me3/H3K9me3 [16–25], collectively regulate luteinization.

Although derived from GCs, luteal cells exhibit a markedly elevated progesterone synthesis capacity, exceeding that of their GC precursors by several orders of magnitude [26]. This metabolic upscaling renders the CL one of the most highly vascularized and energetically demanding tissues. Notably, the intrafollicular microenvironment remains non-vascularized, necessitating *de novo* angiogenesis for efficient substance delivery to luteal cells [27–29]. Paradoxically, vascularization in the developing CL lags behind the increase in progesterone production. This temporal uncoupling of metabolic demand and vascularization creates a transient avascular phase during early luteogenesis, presenting a physiological enigma, i.e., how does the avascular CL meet its high energetic requirements for elevated progesterone production.

This investigation reports a novel cellular adaptive mechanism, GC energy storage (GCES), which addresses this enigma. By using single-cell/spatial transcriptomics, targeted metabolic flux tracing, isotope tracing, and conditional gene silencing, we show that upon luteinization initiation, GCs undergo a coordinated metabolic shift: dampening overall metabolism to reduce energy expenditure coincides with the strategic redirection of absorbed glucose toward glycogenesis, establishing a glycogen storage pool. During the avascular phase, the luteinizing GCs use the stored glycogen to satisfy the energy demanding of elevated progesterone synthesis, thereby ensuring normal luteogenesis. GCES is evolutionarily conserved across species. Disrupting GCES or glycogenolysis precipitate luteal insufficiency, while enhancing GCES improves luteal function and fertility outcomes in mammals.

## RESULTS

### 1. Integrated multi-omics analysis reveals systemic metabolic downregulation GCs during early luteinization

To investigate the cellular response of GCs during the early stage of luteinization, the mouse ovaries harvested at 0, 1 and 6 hours post-hCG (H0, H1 and H6) administration were subjected to single-cell RNA sequencing to capture high-resolution transcriptomes of ovarian cell populations (Figure S1A, B). Complementary spatial transcriptomic analysis of publicly available mouse ovarian datasets [30] enabled construction of a spatially informed transcriptomic atlas of GCs across early luteinization (Figure S1C). By integrating the single-cell and spatial transcriptomic data (Figure 1A; see Materials and Methods), we reconstructed a high-resolution and spatially informed transcriptional atlas of GCs in pre-ovulatory follicles following luteinization initiation (Figure 1B; Table S1). Principal component analysis (PCA) revealed significant transcriptional divergence in GCs across timepoints (Figure 1C), with integrated data demonstrating acceptable quality (gene training scores: ∼0.8; AUC: 0.63–0.70) and enhanced resolution versus original spatial data (Figure 1D-F).

**Figure 1.**
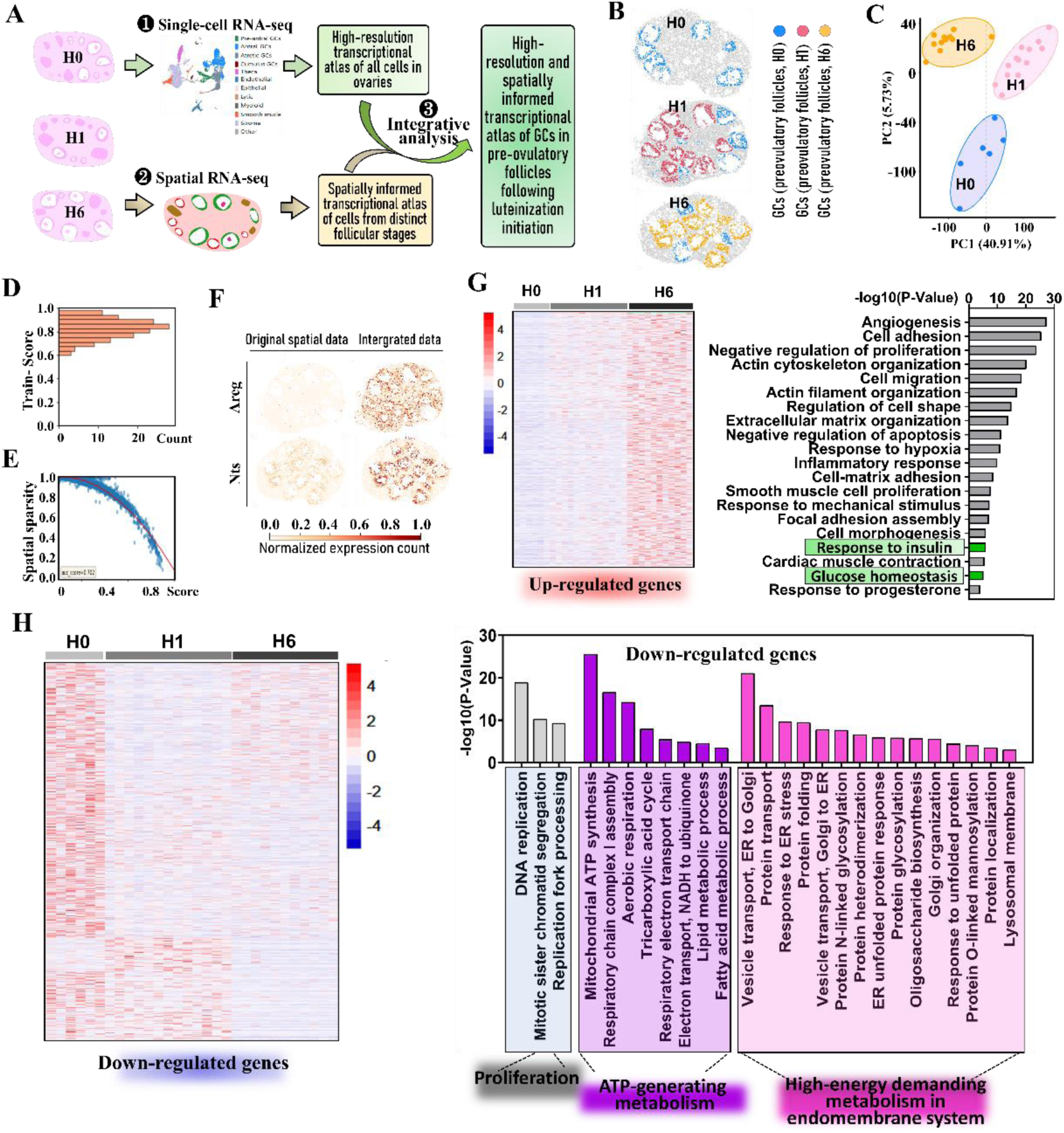
Integrated multi-omics analysis reveals systemic metabolic downregulation GCs during early luteinization. (A) Workflow for integrative analysis of single-cell and spatial transcriptomics data. (B) Identification of GCs in pre-ovulatory follicles at specified time points post-hCG administration using integrated analysis. (C) PCA analysis of transcriptomic differences in GCs from pre-ovulatory follicles. (D) Histogram of gene similarity scores derived from model training data. (E) Scatter plot of tangram-predicted gene scores (x-axis) versus spatial sparsity (y-axis) based on integrated transcriptome data. (F) Expression analysis of randomly selected genes demonstrated enhanced spatial resolution and clearer localization patterns in the integrated dataset compared to the original spatial transcriptomics data. (G) GO enrichment analysis of the upregulated expression cluster. Left: Heatmap of cluster-specific genes; Right: Enriched GO terms. (H) GO enrichment analysis of the downregulated expression cluster. Left: Heatmap of cluster-specific genes; Right: Enriched GO terms.

Differential expression analysis identified 3000 significantly upregulated and 3187 downregulated genes in GCs following luteinization initiation. Gene Ontology (GO) analysis demonstrated that the upregulated genes predominately enriched in established luteinization-associated processes, including *Cell Migration*, *Angiogenesis*, *Cell Adhesion*, *Extracellular Matrix Organization*, *Inflammatory Response* and *etc*, aligning with the observations of previous studies, however, two processes–*Response to Insulin* and *Glucose Homeostasis*–were newly identified (Figure 1G).

Interestingly, downregulated genes exhibited striking enrichment in diverse yet interconnected biological processes: *Mitochondrial ATP Synthesis*, *Respiratory Chain Complex I Assembly*, *Aerobic Respiration*, *Tricarboxylic Acid (TCA) Cycle*, *Respiratory Electron Transport Chain*, *Electron Transport, NADH to Ubiquinone*, *Lipid Metabolic Process*, *Fatty Acid Metabolic Process*, *Vesicle transport, ER to Golgi*, *Response to ER Stress*, *Protein Folding*, *Vesicle Transport, Golgi to ER*, *Protein N-glycan Processing*, *Protein Heterodimerization*, *ER Unfolded Protein Response*, *Protein Glycosylation*, *Oligosaccharide Biosynthesis*, *Golgi Organization*, *Protein O-linked Mannosylation*, *Protein Localization*, *Lysosomal Membrane* (Figure 1H). These processes collectively implicate a metabolic reprogramming toward quiescence, characterized by suppressed ATP-generating activities in mitochondria and energy-demanding activities in the endomembrane system. This transcriptional shift provides preliminary evidence suggesting a hitherto unrecognized cellular adaptation wherein GCs adopt energy-conserving strategies during early luteinization, suppressing ATP-generating and energy-consuming processes.

### 2. Metabolic attenuation and bioenergetic repression in GCs characterize early luteinization

The transcriptomic predictions of bioenergetic suppression were corroborated through multimodal analyses, including ¹³C-glucose tracing, ultrastructural imaging, and bioenergetic profiling. Metabolic flux analysis demonstrated diminished ¹³C-glucose labeling in glycolysis and TCA cycle intermediates following hCG administration (Figure 2A, B; S2A). Ultrastructural remodeling of mitochondria was evident, transitioning from rounded at H0 to elongated morphologies at H6, concomitant with significant reduction in ER-mitochondrial contact sites–critical platforms for metabolic crosstalk (Figure 2C) [31]. This architectural reorganization paralleled profound bioenergetic suppression, characterized by a 27.0% decline in mitochondrial membrane potential (MMP), a 2.90-fold increase in NAD⁺/NADH ratio, and a 44.1% decrease in ATP synthesis relative to H0 controls (Figure 2D-F).

**Figure 2.**
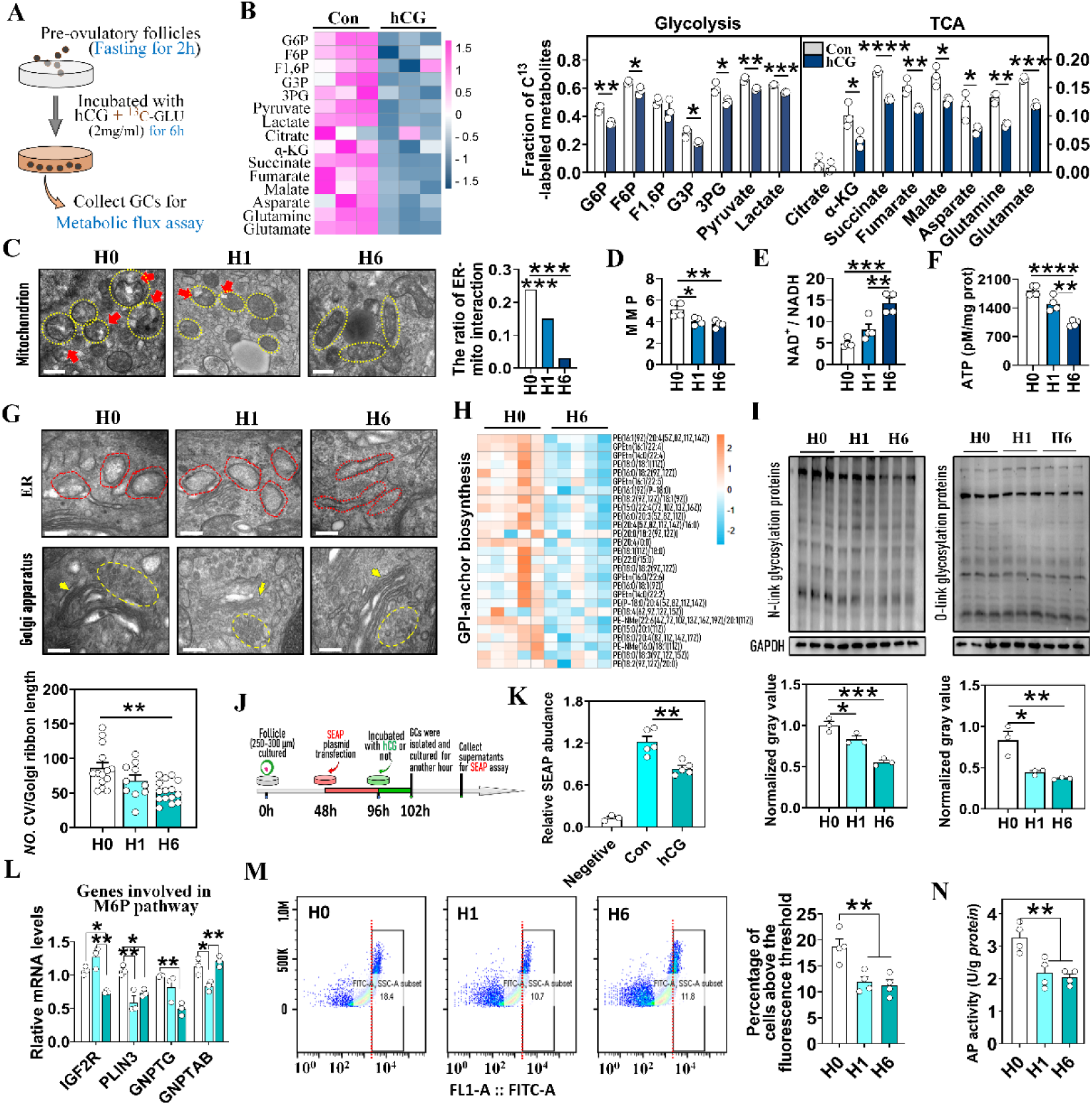
Metabolic attenuation and bioenergetic repression in GCs characterize early luteinization. (A) Workflow for metabolic flux analysis. Pre-ovulatory follicles were fasted for 2 hours, then cultured in luteinization medium with U-¹³C6-glucose as sole carbon source for 6 hours before GC isolation and targeted metabolomics. (B) Quantitative analysis of ¹³C-labeled glycolytic and TCA cycle intermediates. Left: heatmap; Right: quantification (n = 3 GC samples). The control was defined as the group cultured in hCG-free medium. (C) Changes in mitochondrial ultrastructure and ER-mitochondria contact sites following hCG injection. Left: representative images. Right: quantification. Yellow outlines: mitochondria; red arrow: contact sites. Scale bar = 500 nm. (D) MMP measured by JC-1 staining (n = 4 GC samples). (E) NAD⁺/NADH ratio determined by colorimetric assay (n = 4 GC samples). (F) ATP levels measured by luciferase bioluminescence assay (n = 4 GC samples). (G) Ultrastructural changes in ER and Golgi apparatus in GCs following hCG injection. Top: representative images. Down: quantification. Red outlines: ER; yellow arrow: Golgi apparatus; yellow outline: transport vesicles. Scale bar = 200 nm. (H) Heatmap showing changes in the content of GPI-anchor biosynthesis pathway intermediates post-hCG (n = 5 GC samples). (I) Quantification of N-linked (Stained by Concanavalin A)/O-linked glycoprotein (Stained by Peanut Agglutinin) abundance in GCs by lectin blotting. Top: lectin blot; Down: quantification (n = 3 GC samples). (J) Workflow for SEAP reporter assay. (K) Quantification of SEAP secreted into the supernatant using chemiluminescence assay (n = 5 supernatant samples). (L) Changes in M6P pathway gene expression post-hCG injection (n = 3 GC samples). (M) Flow cytometry analysis of GCs with high-intensity Lyso-tracker green fluorescence. Red line: fluorescence intensity threshold; cells right of threshold line classified as high-intensity fluorescent (n = 4 GC samples). (N) Detection of changes in lysosomal acid phosphatase activity using colorimetric assay (n = 4 GC samples). Data represent mean ± SD. Significance determined by two-tailed unpaired Student’s t-test (G) and one-way ANOVA followed by Tukey’s post hoc test (*P<0.05, **P<0.01, ***P<0.001, ****P<0.0001). Results shown (C-G, I-N) are from one of at least three independent replicates; all experiments produced comparable data.

We next validated the attenuation of metabolic activity within the endomembrane system through quantitative assessment of protein glycosylation, vesicular trafficking, secretory capacity, and lysosomal biogenesis. Ultrastructural examination revealed that GCs at H0 exhibited dilated endoplasmic reticulum (ER) cisternae and accumulated transport vesicles adjacent to Golgi apparatus. By 6 hours post-hCG, ER tubules had transformed into elongated cord-like structures, concurrent with a substantial reduction in peri-Golgi vesicular density (Figure 2G). This morphological remodeling correlated with the coordinated downregulation of genes regulating N-linked glycosylation, oligosaccharide biosynthesis, vesicle assembly, and Golgi-specific O-glycosylation (Figure S2B-D). Metabolomic profiling further indicated a general decline in intermediates of the glycosylphosphatidylinositol (GPI) anchor biosynthesis pathway–critical for glycosylation–in H6 GCs (Figure 2H), a finding corroborated by lectin binding assays demonstrating significantly reduced abundance of glycosylated proteins post-hCG (Figure 2I; S2E). Quantitative assessment of secretory capacity via secreted alkaline phosphatase (SEAP) reporter transfection demonstrated a significant reduction in SEAP activity post-hCG (Figure 2J, K; S2F). Furthermore, genes in the mannose-6-phosphate (M6P) pathway—essential for lysosomal biogenesis—were significantly downregulated post-hCG (Figure 2L), aligning with reduced LysoTracker Green fluorescence intensity (Figure 2M) and decreased lysosomal acid phosphatase activity (Figure 2N).

These data demonstrate metabolic quiescence transition in GCs during early luteinization, validating the omics predictions.

### 3. Enhanced glucose uptake supports glycogen biosynthesis in GCs during early luteinization

Intriguingly, despite transitioning to a metabolically quiescent state, GCs at H6 exhibited a significant increase in glucose uptake compared to H0 controls (Figure 3A). Transporter expression profiling identified SLC2A1 as the predominant glucose transporter of this enhanced uptake (Figure 3B; S3A). Intracellular glucose can be directed toward ATP generation or storage as lipids or glycogen (Figure 3C) [32]. Metabolic flux analysis ruled out significant catabolic utilization (Figure 2A, B), and nor for the lipid storage since colorimetric assays revealed the reduced intracellular free fatty acids and triglycerides (Figure S3B, C). To directly assess glycogen conversion, we performed ¹³C-glucose tracing coupled with Liquid Chromatograph-Tandem Mass Spectrometer (LC-MS/MS). hCG-treated GCs exhibited pronounced ¹³C enrichment in glycogen (35% incorporation versus, 8% in controls; Figure 3D, E), demonstrating preferential glycogen biosynthesis. Consistent with this, Western blotting and qRT-PCR analysis detected activation of the glycogenesis pathway following hCG administration (Figure 3F; S3D). Complementary Periodic Acid-Schiff (PAS) staining and colorimetric quantification confirmed a significant elevation in glycogen content within H6 GCs (Figure 3G, H).

**Figure 3.**
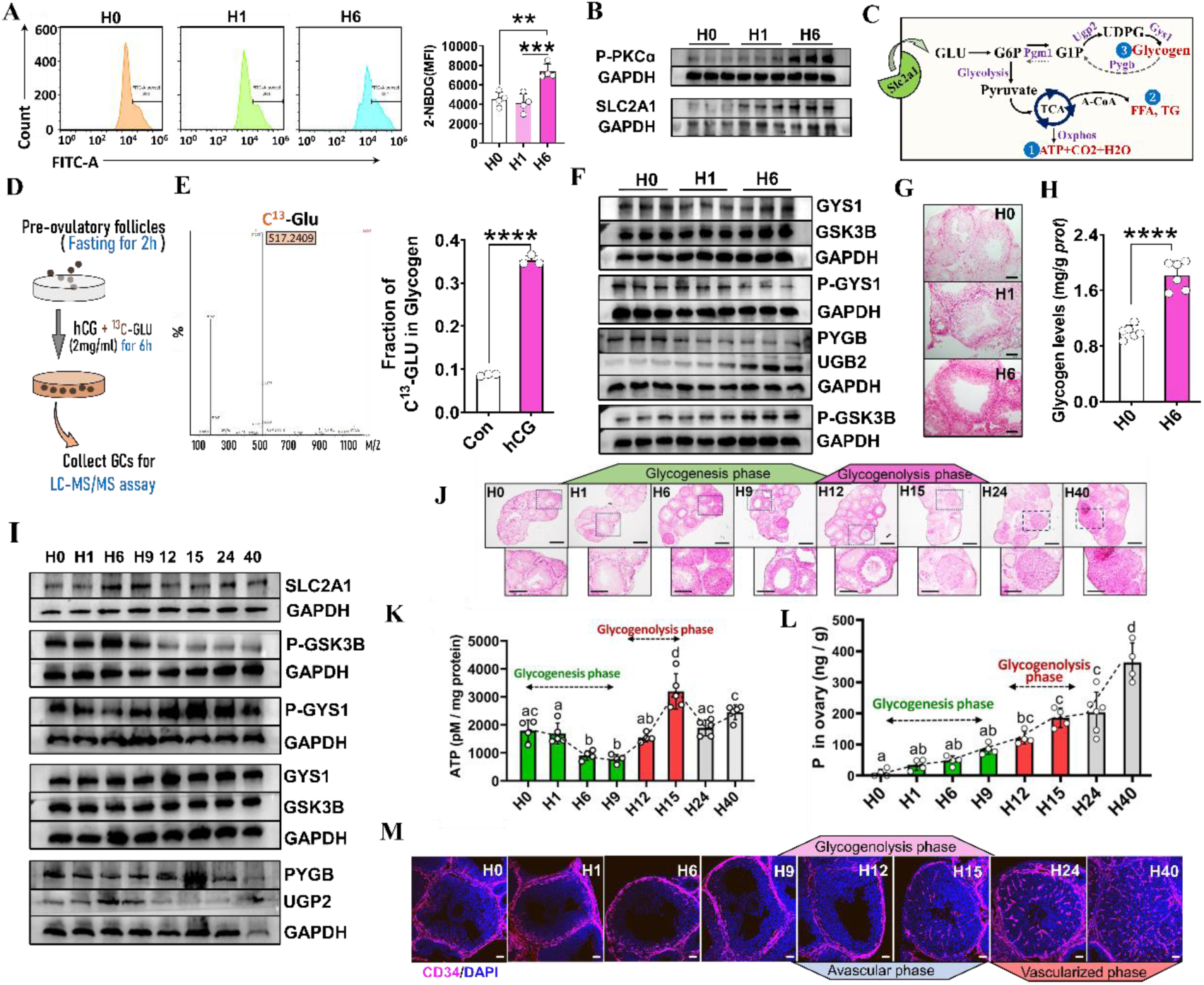
Enhanced glucose uptake supports glycogen biosynthesis in GCs during early luteinization. (A) Flow cytometric analysis of cellular glucose uptake capacity post-hCG injection. Isolated GCs were incubated with fluorescent analog 2-NBDG (100 μM) for 30 minutes prior to analysis. Left: representative images; Right: quantification (n = 4 GC samples). (B) Western blotting analysis of SLC2A1 and its upstream kinase P-PKCα in GCs post-hCG injection (n = 3 GC samples). (C) Schematic of intracellular glucose carbon flux. (D) ^13^C-glucose tracing workflow. Pre-ovulatory follicles fasted 2 hours, then cultured in luteinization medium with U-¹³C6-glucose (sole carbon source) for 6 hours before GC isolation and LC-MS/MS analysis. (E) Quantitative ¹³C enrichment in glycogen (n = 3 GC samples). The control was defined as the group cultured in hCG-free medium. (F) Changes in glycogenesis/glycogenolysis proteins during early luteinization (n = 3 GC samples). (G) PAS staining of glycogen in GCs during early luteinization. Scale bar = 100 μm. (H) Colorimetric quantification of glycogen content (n = 6 GC samples). (I) Changes in glycogenesis/glycogenolysis proteins throughout luteinization. (J) PAS staining of glycogen throughout luteinization. Scale bar = 500 μm (high-magnification), 250 μm (low-magnification). (K) ATP content changes throughout luteinization (n = 5-6 GC samples). (L) Radioimmunoassay of progesterone in ovarian homogenates throughout luteinization (n = 4-7 ovarian samples). (M) Vascularization changes assessed by CD34⁺ endothelial cells (red) during luteinization. Nuclei counterstained with DAPI (blue). Scale bar = 50 μm. Data represent mean ± SD. Significance determined by one-way ANOVA with Tukey’s post hoc test (A, K, L) or two-tailed unpaired Student’s t-test (E, H) (*P < 0.05, **P < 0.01, ***P < 0.001, ****P < 0.0001). Results shown are from one of at least three independent replicates; all experiments produced comparable data.

We next characterized the temporal dynamics of glycogen storage and its functional correlates throughout luteinization process. Integrated analysis defined distinct phases: a glycogenesis phase (H0-H9) followed by a glycogenolysis phase (H9-H15), evidenced by glycogenesis/glycogenolysis pathway protein expression (Figure 3I) and PAS staining (Figure 3J). Notably, during glycogenesis phase, ATP production declined (Figure 3K) while progesterone levels rose gradually up to 80 ng/g (Figure 3L). Upon entering glycogenolysis phase (H9-H15), ATP levels surged (Figure 3K) coincident with the peak progesterone secretion (∼200 ng/g; Figure 3L). Critically, this glycogenolysis phase–characterized by high energy demand for steroidogenesis–occurred within the developing CLs prior to the establishment of internal vascularization (Figure 3M).

These data demonstrate that during the early luteinization phase, GCs undergo a coordinated metabolic shift: dampening overall metabolism to reduce energy expenditure coincides with the strategic redirection of absorbed glucose toward glycogenesis, establishing a glycogen storage pool. We postulate that this preformed glycogen serves as a critical energy reservoir for the avascular CL.

### 4. Mobilization of glycogen storage to fuel luteal steroidogenesis during the avascular phase

To test whether the stored glycogen in GCs is used to fuel the avascular CL for steroidogenesis, we augmented glycogen storage during glycogenesis phase via 2g/kg dose of glucose (glucose tolerance dose) administration (Figure 4A-C). By 15 hours post-hCG (H15), the glucose-loaded luteinizing GCs exhibited reduced NAD⁺/NADH ratio, increased ATP generation, upregulated luteal gene expression and increased progesterone secretion compared to the controls (Figure 4D, E; S4A). This functional enhancement persisted until H40 (Figure S4B), indicating the glycogen storage as a critical determinant of luteal function.

**Figure 4.**
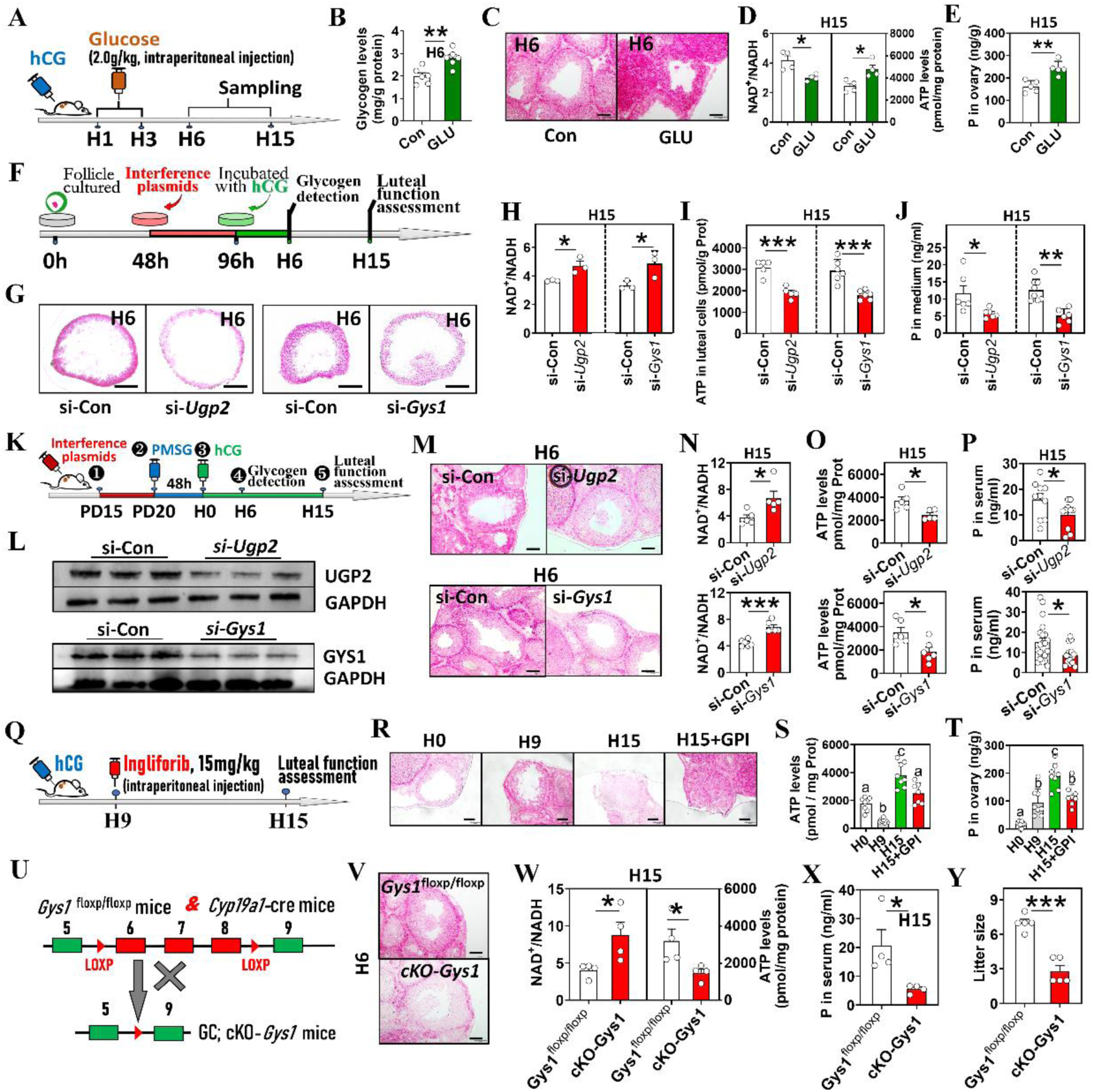
Mobilization of glycogen storage to fuel luteal steroidogenesis during the avascular phase. (A) Experimental design for panels (B-E). (B) Colorimetric quantification of glycogen content in GC post-glucose injection (n = 6 GC samples). (C) PAS staining of glycogen storage post-glucose injection; scale bar = 100 μm. (D) Alterations in NAD⁺/NADH ratio and ATP contents post-glucose injection (n = 4 GC samples). (E) Progesterone levels in ovarian homogenates measured by radioimmunoassay post-glucose injection (n = 5 ovarian samples). (F) Experimental design for *Ugp2*/*Gys1* knockdown in cultured mouse follicles. Luteinizing GCs isolated at H15 for functional assessment. (G) Changes in GC glycogen storage post-*Ugp2* and *Gys1* knockdown. The commercial scrambled siRNA serving as si-Control (si-Con). Scale bar = 100 μm. (H) Alterations in NAD^+^/NADH in GCs post-knockdown; n = 3 GC samples. (I) ATP content in GCs post-knockdown (n = 5-6 GC samples). (J) Progesterone in culture medium post-knockdown (n = 6-8 samples). (K) Experimental design for in vivo lentiviral-mediated *Ugp2*/*Gys1* knockdown. Luteinizing GCs isolated from CLs at H15 for functional assessment. (L) Western blotting confirmation of *Ugp2*/*Gys1* knockdown in ovaries (n = 3 ovarian samples). (M) PAS staining of glycogen storage post-knockdown. Scale bar = 100 μm. (N) NAD⁺/NADH alterations in GCs post-knockdown (n = 5-6 GC samples). (O) ATP content changes in GCs post-knockdown (n = 5-6 GC samples). (P) Progesterone level alterations post-knockdown; n = 10 (si-*Ugp2*), 15-21 (si-*Gys1*) serum samples. (Q) Experimental design for Ingliforib treatment in vivo. (R) PAS staining validation of Ingliforib-mediated glycogenolysis inhibition. Scale bar = 100 μm. (S) ATP levels in luteinizing GCs post-Ingliforib treatment (n = 7-8 GC samples). (T) Progesterone in ovarian homogenates post-Ingliforib treatment (n = 8 ovarian samples). (U) Schematic representation of the conditional knockout of *Gys1* in mouse GCs. (V) PAS staining of glycogen storage in GCs post-knockout. Scale bar = 100 μm. (W) Alterations in NAD⁺/NADH ratio and ATP contents (n = 4 GC samples). (X) Progesterone level alterations post-knockout (n = 4 serum samples). (Y) Litter size alterations post-knockout (n = 5 mice). Data represent mean ± SD. Significance determined by two-tailed unpaired Student’s t-test or one-way ANOVA with Tukey’s post hoc test (S, T) (*P < 0.05, **P < 0.01, ***P < 0.001). Results shown are from one of twice (V-Y) or at least three (A-T) independent replicates; all experiments produced comparable data.

To further validate the functional importance of glycogen, RNA interference was performed to suppress the glycogen synthesis. *In vitro* knockdown of glycogenesis genes *Ugp2* and *Gys1* did not impair follicular development and ovulation but markedly reduced GC glycogen storage in pre-ovulatory follicles (Figure 4F, G; S5A-C). This intervention precipitated a profound bioenergetic crisis at H15, characterized by increased NAD⁺/NADH ratio, diminished MMP, reduced ATP generation, significantly suppressed progesterone synthesis and luteal gene expression (Figure 4H-J; S5D, E). *In vivo* ovarian bursa delivery of *Ugp2* and *Gys1* interference plasmids (Figure 4K-M; S6A) recapitulated this bioenergetic crisis with: increased NAD⁺/NADH, decreased ATP, downregulated luteal gene expression and reduced serum progesterone (Figure 4N-P; S6B). Complementarily, inhibition of glycogen catabolism during the glycogenolysis phase using the PYGB inhibitor–Ingliforib (Figure 4Q, R) also induced luteal insufficiency, evidenced by decreased ATP production, downregulated luteal gene expression and attenuated progesterone synthesis in the glycogen-constrained luteinizing GCs (Figure 4S, T; S6C).

To definitively investigate the functional necessity of glycogen storage *in vivo*, the GC-specific *Gys1* conditional knockout mice (cKO-*Gys1*) were generated using a Gyp19a1-Cre driver system (Figure 4U; S6D). This genetic ablation abolished glycogen accumulation within GCs of the mice (Figure 4V). Concomitantly, cKO mice exhibited significant bioenergetic perturbation in their avascular CLs, including elevated NAD⁺/NADH ratio, impaired ATP production and significantly suppressed progesterone synthesis along with attenuated luteal gene expression at H15 (Figure 4W, X; S6E). Moreover, cKO mice displayed a significant decrease in litter size compared to *Gys1*^floxp/floxp^ (Figure 4Y).

These data confirm that glycogenesis in GCs establishes an essential energy reservoir, which mobilized during the avascular phase to fuel the bioenergetic demands of progesterone production in nascent CL. This cellular adaptive mechanism hereafter termed GCES.

### 5. hCG (LH) induces glycogen storage via MAPK-RUNX1-Insulin signaling axis

To identify the intracellular signaling regulating GCES, KEGG enrichment analysis was performed. The results showed a significant induction of insulin signaling during early luteinization (Figure 5A). Consistent with this, qRT-PCR and Western blotting demonstrated that hCG injection markedly increased both mRNA and protein levels of insulin/IGF-1 receptors (INSR and IGF1R) in GCs, concomitant with enhanced receptor phosphorylation (Figure 5B; S7A). Functional validation demonstrated that insulin alone was sufficient to enhance glucose uptake and glycogen storage in GCs. Conversely, pharmacological inhibition of INSR/IGF1R with BMS536924 abolished hCG-induced glucose uptake and glycogen storage (Figure 5C-F), confirming the necessity of insulin signaling for hCG-driven glycogen storage.

**Figure 5.**
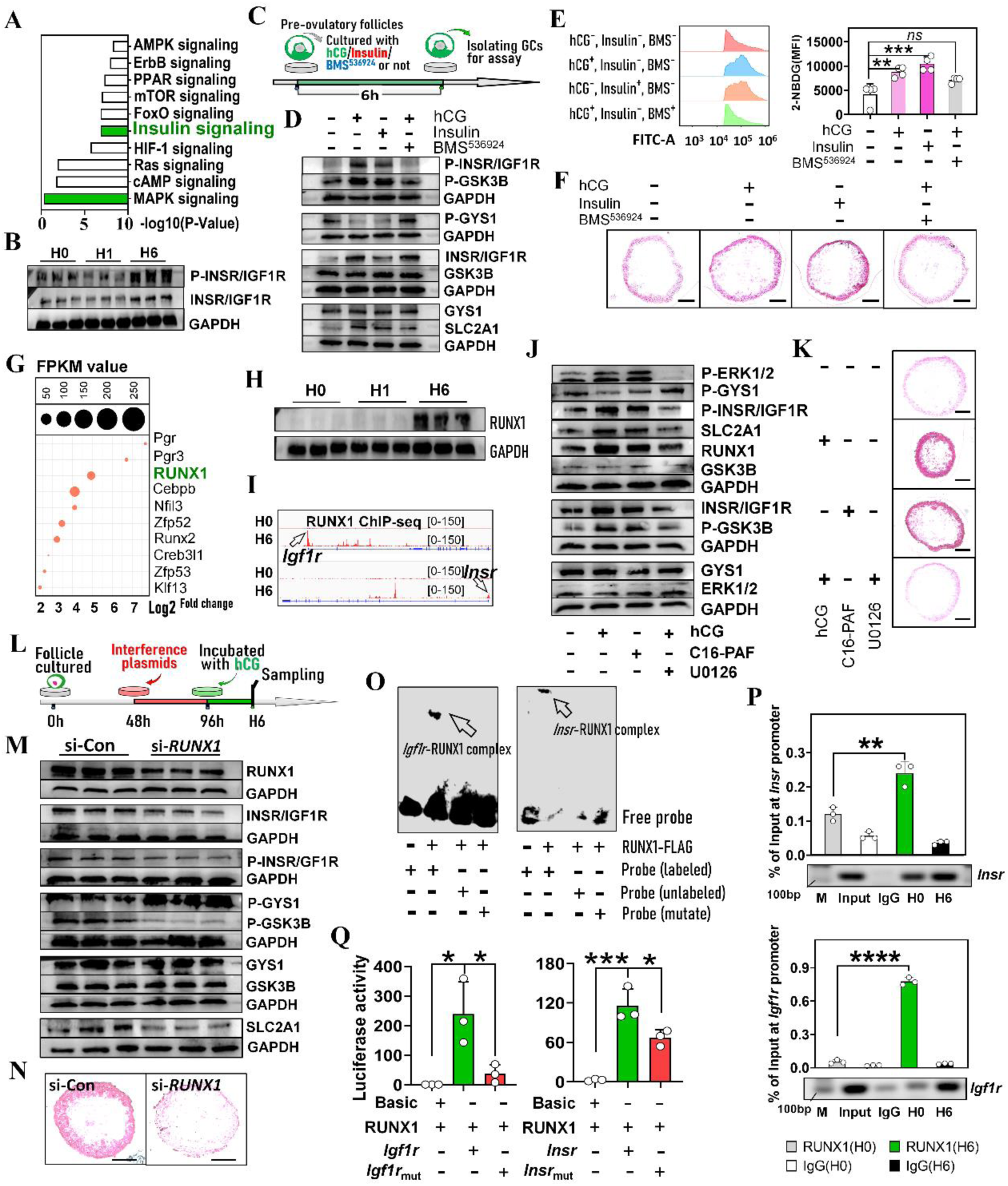
hCG (LH) induces glycogen storage via MAPK-RUNX1-Insulin signaling axis. (A) KEGG pathway analysis of signaling activation in GCs post-hCG injection. (B) Western blotting analysis of INSR/IGF1R activity in GCs post-hCG injection (n = 3 GC samples). (C) Experimental design for panels (D-F). (D) Western blotting analysis of glycogenesis related proteins in GCs after insulin pathway modulation. (E) Flow cytometric analysis of glucose uptake capacity in GCs. Left: representative images; Right: quantification. Isolated GCs were incubated with fluorescent analog 2-NBDG for 30 minutes prior to analysis (n = 4 GC samples). (F) PAS staining of glycogen stores in GCs after insulin signaling manipulation. Scale bar = 100 μm. (G) Top 10 transcription factors upregulated in GCs by hCG (bubble plot). (H) Western blotting analysis of RUNX1 expression in GCs post-hCG injection (n = 3 GC samples). (I) ChIP-seq profiles of RUNX1 binding at *Igf1r* and *Insr* promoters. Arrows: enrichment peaks. (J) Western blotting analysis of RUNX1 and glycogenesis related proteins in GCs after MAPK pathway modulation. (K) PAS staining of glycogen stores after MAPK pathway modulation. Scale bar = 100 μm. (L) Experimental design for panels (M, N). (M) Western blotting analysis of glycogenesis regulators in GCs post-RUNX1 knockdown (n = 3 GC samples). (N) PAS staining of glycogen stores after RUNX1 knockdown. Scale bar = 100 μm. (O) EMSA validating RUNX1 binding to *Igf1r* and *Insr* promoter motifs. (P) ChIP-qPCR quantification of RUNX1 binding to *Igf1r* and *Insr* promoters (n = 3 GC samples). (Q) Dual-luciferase reporter assays identifying RUNX1-responsive elements in *Igf1r* and *Insr* promoters (n = 3 cellular samples). Data represent mean ± SD. Significance determined by one-way ANOVA with Tukey’s post hoc test (*P < 0.05, **P < 0.01, ***P < 0.001, ****P < 0.0001). Results shown (B-F, H, J-Q) are from one of at least three independent replicates; all experiments produced comparable data.

We next investigated mechanism by which hCG (LH) initiates insulin signaling. In addition to insulin signaling, the significant induction of the MAPK (ERK1/2) signaling during early luteinization was also observed with KEGG analysis (Figure 5A). Among the transcription factors potently induced by hCG, RUNX1–a known downstream target of MAPK–was ranked relatively higher (Figure 5G, H). Moreover, publicly available ChIP-seq data [23] indicated specific binding of RUNX1 to *Igf1r* and *Insr* promoters (Figure 5I). Taken together, these data collectively suggest that hCG activates insulin signaling and glycogenesis via the MAPK-RUNX1 cascade. To test this, the C16-PAF was used to active MAPK signaling. This treatment recapitulated the effects of hCG supplementation including RUNX1 upregulation, insulin signaling activation and glycogen storage in GCs. Conversely, MAPK inhibition with U0126 abolished all these effects induced by hCG (Figure 5J, K). Furthermore, RUNX1 knockdown in cultured follicles suppressed hCG-induced insulin signaling activation and glycogen storage in GCs (Figures 5L-M; S7B).

Finally, to identify whether there is a direct transcriptional regulation between RUNX1 and insulin/IGF-1 receptors, both electrophoretic mobility shift assays (EMSA) and chromatin immunoprecipitation quantitative PCR (ChIP-qPCR) were performed. EMSA confirmed the specific RUNX1 binding sites with the predicted consensus motifs within the *Igf1r* and *Insr* promoters (Figure 5O; S7C-E). ChIP-qPCR confirmed an increased RUNX1 occupancy at these promoter regions at H6 (Figure 5P). Site-directed mutagenesis localized the RUNX1 response element to the TGTGGT motif in both promoters (Figures 5M; S7F). These results demonstrate that hCG (LH)-MAPK-RUNX1-Insulin signaling pathway is a key regulator of GCES.

### 6. GCES augmentation enhanced luteal function and pregnancy outcomes in mouse and ovine models

To assess the clinical relevance of GCES, glycogen storage in mouse GCs was augmented with timely glucose administration (Figures 6A, B). This intervention significantly elevated progesterone levels by gestation day 15 (GD15; Figure 6C). Crucially, the progesterone-enhancing effect of glucose administration was contingent exclusively into the glycogenesis phase; glucose administration outside this temporal window was proved no effective (Figure S8A). Compared to controls, the glucose-treated mice exhibited a reduced fetal regression rate (Figure 6D), increased fetal and placental weights (Figure 6E), although fetal number remained unchanged (Figure 6D). To critically evaluate the functional relevance to assisted reproduction, we further assessed the impact of glucose administration on the farrowing rate following embryo transfer (Figure 6F). Glucose treatment during the temporal window significantly increased the farrowing rate compared to the control group (Figure 6G), demonstrating its potential utility in reproductive technologies.

**Figure 6.**
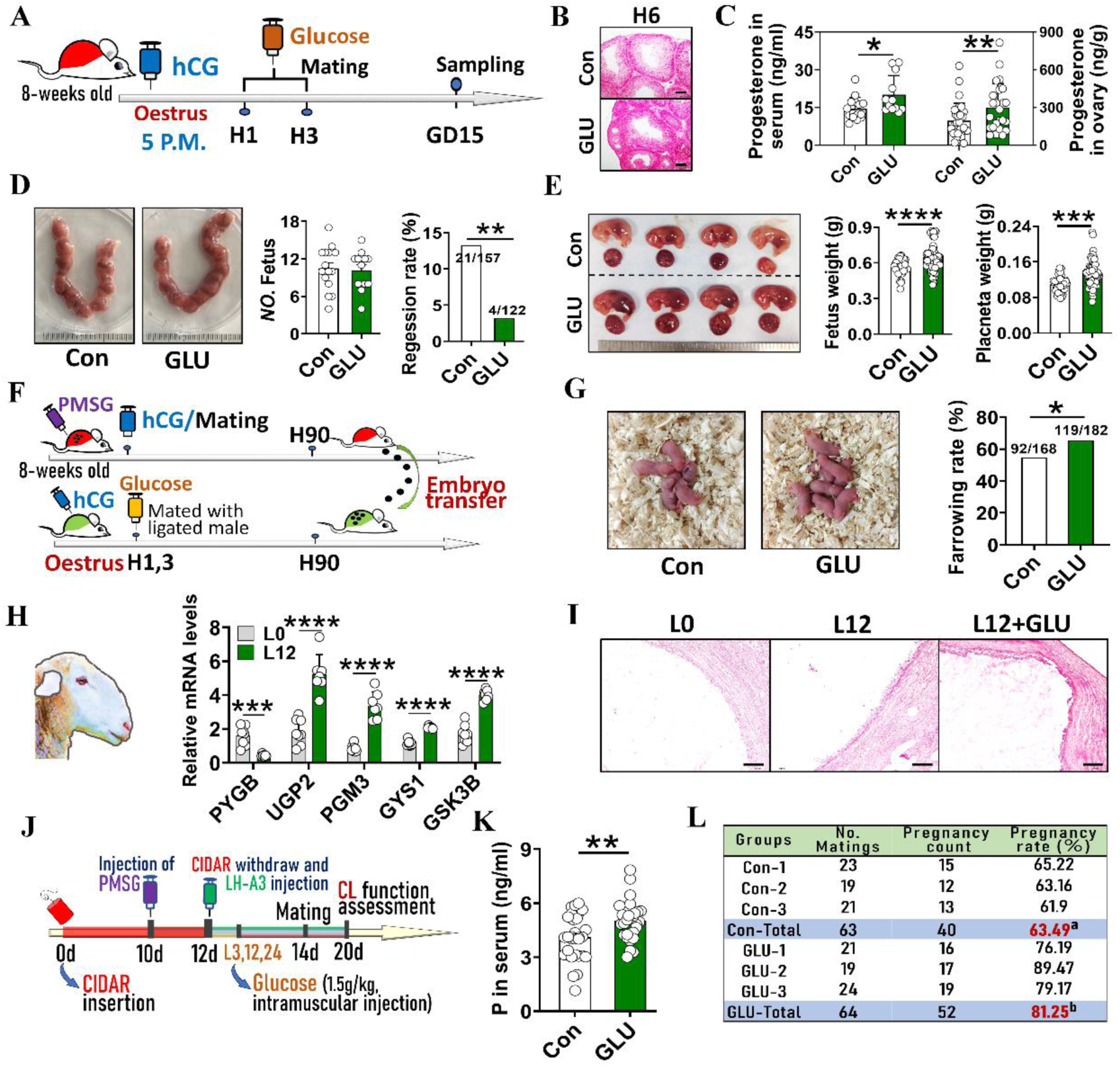
GCES augmentation enhanced luteal function and pregnancy outcomes in mouse and ovine models. (A) Experimental design for panels B to E. (B) PAS staining of glycogen. Scale bar = 100 μm. (C) Effects of glucose intake on serum and ovarian progesterone levels at GD15; n (serum) = 15 (Con), 12 (GLU); n (ovarian homogenate) = 30 (Con), 24 (GLU). (D) Effects of glucose intake on fetal number and regression rates at GD15 (n = 15 pregnant mice). (E) Effects of glucose intake on fetal and placental weights at GD15; n = 4 (Con) and 5 (GLU) pregnant mice. (F) Embryo transfer experimental schematic. (G) Effects of glucose intake on the farrowing rate of transferred embryos. (H) qRT-PCR analysis of glycogen metabolism genes in ovine GCs pre-(L0) and 12 hours post-LH injection (L12); n = 8 (L0), 7 (L12) GC samples. (I) PAS staining of glycogen in ovine GCs post-LH injection. Scale bar = 500 μm. (J) Experimental design for panels I to L. (K) Effects of glucose intake on serum progesterone levels (n = 30 serum samples). (L) Effects of glucose intake on sheep pregnancy rate after breeding. Data represent mean ± SD. Significance determined by two-tailed unpaired Student’s t-test or chi-square test (D, G and L) (*P < 0.05, **P < 0.01, ***P < 0.001, ****P < 0.0001). Results shown are from one of at least three independent replicates; all experiments produced comparable data.

The functional conservation of GCES was also investigated in sheep, a mono-ovulatory species. Similar to the mouse model, LH injection significantly upregulated the expression of genes involved in insulin and glycogenesis pathway, reduced ATP generation and increased glycogen accumulation in sheep GCs (Figures 6H, I; S8B, C). The subsequent glucose potentiation (1.5g/kg, aligned with human glucose tolerance dose) further increased glycogen storage (Figures 6I, J), concomitantly elevating progesterone levels and luteal gene expression by GD20 (Figure 6K; S8D). Strikingly, the pregnancy rate was significantly higher in the glycogen-augmented group (81.25%) compared to controls (63.45%; p < 0.01, Figure 6L).

### 7. Timely oral glucose administration enhances GCES and luteal function in humans

Transitioning to humans, the transcriptome of human GCs following hCG administration was analyzed by using public databases [33]. GO analysis revealed the enrichment of these hCG-downregulated genes in high-energy-demanding processes and ATP production, similar to the findings in mouse and ovine models (Figure 7A). Moreover, the expression of glucose transporters *SLC2A13*, *SLC2A14* and *SLC2A3* exhibited a sharp increase at H12, returning to baseline by H32 (Figure 7B). Furthermore, hCG administration significantly upregulated insulin signaling components, including *IGF1R*, *IGFBP1*, *IGFBP3*, *IGFBP5* and *IRS2* (Figure 7C), supporting the evolutionary conservation of GCES mechanisms in humans.

**Figure 7.**
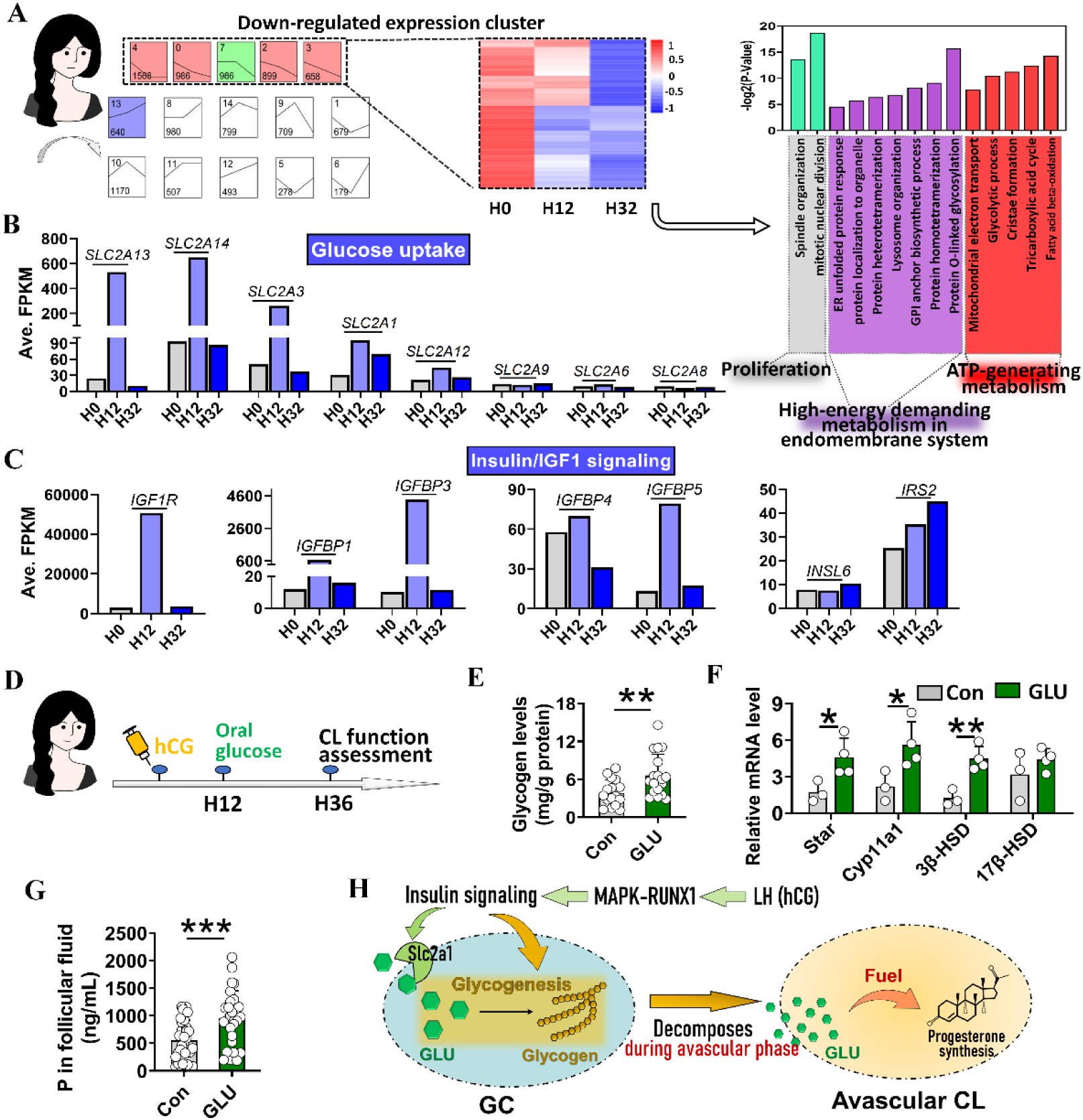
Timely oral glucose administration enhances GCES and luteal function in humans. (A) GO enrichment analysis of the downregulated expression cluster in human GCs post-hCG injection. Left: heatmap; Right: enriched GO terms. (B) Expression of glucose transporter genes in GCs post-hCG injection. (C) Expression of insulin pathway genes in GCs post-hCG injection. (D) Experimental design for panels E to G. Volunteers were randomly selected from women undergoing ART treatment and responding normally to controlled ovarian hyperstimulation. (E) Colorimetric qualification of glycogen storage in GCs post-glucose intake; n = 16 (Con), 18 (GLU) GC samples. (F) qRT-PCR analysis of luteal gene expression post-glucose intake; n = 3 (Con), 4 (GLU) GC samples. (G) Progesterone levels in follicular fluids measured by radioimmunoassay post-glucose intake; n = 34 (Con), 28 (GLU) samples. (H) A diagram depicting GCES. Data represent mean ± SD. Significance determined by two-tailed unpaired Student’s t-test (*P < 0.05, **P < 0.01, ***P < 0.001). The experiments (E, G) were repeated twice with consistent results.

To translate these findings clinically, we recruited volunteers undergoing assisted reproductive technology (ART). Participants were instructed orally intake of 75g glucose (glucose tolerance dose) at 12 hours post-hCG trigger. Their progesterone synthetic capacity was assessed 24 hours post-glucose ingestion (Figure 7D). This timely oral glucose administration significantly augmented glycogen storage within human GCs (Figure 7E). Compared to the control group, the glucose-treated group demonstrated significantly upregulated expression of luteal genes in luteinizing GCs (Figure 7F), culminating in a ∼73% increase (p < 0.001) in mean progesterone concentration within follicular fluid (Figure 7G).

These results demonstrate that a single, timely oral glucose intervention following hCG trigger significantly elevates glycogen reserves in human GCs and increases progesterone production.

## DISCUSSION

The acquisition of progesterone synthesis capacity precedes the vascularization in the CL. How does the avascular CL meet the substantial bioenergetic demands for the high level of progesterone production is still unknown. Our study resolves this enigma by identifying GCES as the key mechanism. We demonstrate that during early luteinization, hCG (LH) orchestrates a metabolic transition in GCs via the MAPK-RUNX1-Insulin signaling axis, redirecting glucose toward glycogenesis. This strategically stored glycogen is subsequently mobilized to fuel intensive progesterone production throughout the avascular phase, ensuring normal luteogenesis (Figure 7H). The GCES process fundamentally refines our understanding of luteinization. This process strikingly parallel to oocyte nutrient provisioning for preimplantation embryos [34] highlights conserved regulatory strategies in reproduction. Moreover, GCs emerge as a novel insulin-sensitive glycogenic cell type, distinct from hepatocytes and myocytes [35,36], with broad physiological implications.

GCES not only serves as an energy source for avascular CL but also profoundly influencing mature luteal function (Figure S4B), indicating significant translational promise. GCES enhancement improved luteal function in both polytocous (mice) and monotocous (sheep) species, leading to marked increases in pregnancy rates. In human trials, the timely oral glucose administration induces the GCES enhancement, increasing progesterone content by ∼73%. Crucially, this GCES-mediated luteal support was achieved solely through glucose intake, bypassing pharmaceutical agents, thereby being exceptional safety. This is particularly significant since no treatment for luteal insufficiency has been reported to improve pregnancy rates in natural menstrual cycle [3]. Therefore, we propose a novel preconception strategy: peri-ovulatory intake of glucose or high-glycemic-index foods (such as cakes, milk tea, candies) as a potential means to mitigate luteal insufficiency and improve fertility outcomes. Given its safety, accessibility, ease of use and cost-effectiveness, this approach could be widely adopted in populations with a higher incidence of luteal insufficiency, potentially reducing reliance on conventional luteal phase support–especially in women who experience the dual risk factors of early pregnancy bleeding and a history of previous miscarriages [37,38].

In livestock production, GCES-based luteal enhancement could integrate seamlessly with estrus synchronization and embryo transfer to boost pregnancy rates and reduce infertility or miscarriage attributable to luteal deficiency, as validated in sheep (Figure 6K, L; S8D). Importantly, optimal implementation requires precision timing, dosage (the dose employed herein was glucose tolerance dose), and frequency of glucose administration. Murine data identified a ≤9-hour post-hCG window (marked by glucose transporter/glycogen synthase upregulation) for effective glucose intervention. Extra-window supplementation fails to improve luteal function (Figure S8A), necessitating species-tailored glucose regimens–a key challenge in clinical translation.

While GCs and theca cells jointly contribute to luteogenesis, GO analysis revealed a distinct metabolic divergence. Unlike GCs, theca cells neither enter metabolic quiesce nor accumulate glycogen following luteinization initiation (Figure S9A-D). Thus, glycogen storage represents a GC-specific adaptation. Intriguingly, theca cells exhibit significantly higher glycogen content compared to other follicular cell types during folliculogenesis (Figure S9E), suggesting that theca cells may also engage in glycogen storage, albeit initiating this process earlier than GCs.

In summary, this study introduces the novel concept of GCES and demonstrates that enhancing GCES via timely glucose administration improves luteal function in different species including humans. These insights advance reproductive physiology and hold substantial promise for developing novel therapeutic approaches to address luteal insufficiency in both livestock and human.

## LIMITATIONS

While we have identified the LH (hCG)-MAPK-RUNX1-Insulin axis regulating glycogen storage in GCs, the signaling pathway(s) inducing metabolic quiescence remains undefined. Beyond elevated energy demands, robust progesterone production also requires substantial cholesterol precursor, yet how the developing CL maintains cholesterol supply during its avascular phase has not been investigated. Furthermore, the mandatory progesterone supplementation following embryo transfer in ART protocols [39,40] precludes direct assessment of GCES-mediated improvement in mature CL function and pregnancy outcomes of humans. Future studies should aim to: (i) delineate the mechanisms governing GC metabolic quiescence and cholesterol provision, and (ii) quantify the effects of GCES enhancement on human pregnancy rates in natural conception cohorts.

## MATERIALS AND METHODS

### Experimental design

This study commenced with the prediction of GC metabolic alterations during early luteinization using multi-omics profiling. These predictions were experimentally validated, defining a novel cellular adaptive mechanism, GCES. The functional impact of GCES on luteogenesis was then investigated through targeted manipulation in vitro and in vivo. Meanwhile, the regulatory pathway controlling GCES was elucidated. Finally, the translational potential of GCES was evaluated across species and in a preliminary clinical setting.

1. **Multi-Omics Profiling of GC Dynamics during Early Luteinization**: Integrated single-cell and spatial transcriptomics analysis was performed to delineate high-resolution transcriptional profiles of GCs within pre-ovulatory follicles at 0, 1, and 6 hours post-hCG administration, simulating the LH surge. Subsequent GO enrichment analysis of transcriptomic profiles predicted metabolic alterations in GCs during early luteinization, revealing attenuated high-energy demanding and ATP-generating activities alongside enhanced insulin responsiveness and glucose homeostasis.
2. **Experimental Validation of Metabolic Alterations**: Next, a multi-modal approach was employed to validate the omics-derived predictions, encompassing: targeted metabolomics, electron microscopy, flow cytometry, mitochondrial functional assays, glucose uptake capacity measurements, isotopic tracing, qRT-PCR, Western blotting, immunofluorescence, PAS staining, and enzymatic activity assays. This validation confirmed that GCs enter a metabolically quiescent state while concurrently enhancing glucose uptake and conversion to glycogen for storage, *i.e.* GCES.
3. **Functional Impact of GCES on luteinization**: To investigate whether GCES governs luteal function by modulating energy availability during the avascular phase, glycogen storage was manipulated assessed both in vitro and in vivo via: (i) glucose supplementation to enhance glycogen storage; (ii) RNA interference and conditional knockout strategies to block glycogen storage; and (iii) pharmacological inhibition of glycogen catabolism. The functional consequences of these interventions on luteal energy supply and progesterone biosynthesis were rigorously evaluated.
4. **Validation of GCES Regulatory Pathway**: Bioinformatics-driven analyses, coupled with flow cytometry, pharmacological modulation (inhibitors/activators), RNA interference, Western blotting, EMSA, ChIP-qPCR, luciferase reporter assay, and PAS staining, were utilized to elucidate the signaling pathway(s) governing GCES.
5. **Translational Evaluation of GCES in a Three-Phase Approach**: (i) The conservation of the GCES phenomenon was validated across species (mouse, sheep, human); (ii) The impact of augmenting GCES via post-hCG glucose administration on luteal function, pregnancy rate, and embryo transfer success was systematically evaluated in mouse/ovine models; (iii) Volunteers undergoing ART treatment were recruited. Following hCG administration, participants received an oral glucose administration at 12 hours to augment GCES. The efficacy of this intervention in enhancing early human luteal function was assessed.

### Animals

KM strain mice were sourced from the Center for Animal Testing at Huazhong Agricultural University (Wuhan, China), while *Gys1*^floxp/floxp^ C57BL/6J mice [41] were obtained from GemPharmatech (Nanjing, China). GC-specific deletion of *Gys1* exons 6-8 was achieved by crossing *Gys1*^floxp/floxp^ mice with Cyp19a1-Cre transgenic mice [42] (Cyagen; Suzhou, China). All mice were maintained under specific pathogen-free (SPF) conditions with controlled temperature (25 ± 1°C), a 12-hour light/dark cycle, and ad libitum access to food and water. Sheep subjects were housed at Xinjiang Jinken Aoqun Agriculture and Animal Husbandry Technology (Hetian, China) following standard husbandry protocols. Experimental procedures strictly adhered to the institutional guidelines for laboratory animal welfare and were approved by the Animal Ethics Committee of Huazhong Agricultural University (Approval No.: HZAUMO-2023-0115; HZAUSH-2024-0001).

### Human clinical trials

Female participants (25-40 years) undergoing ART treatment were prospectively enrolled in this clinical study. Following comprehensive verbal and written disclosure of study objectives, procedures, and potential risks/benefits, written informed consent was obtained from all subjects. At 12 hours post-hCG trigger (Ovidrel®, 250 μg; Sigma, USA), participants were instructed to ingest 75g glucose (Chinaotsuka, China). Transvaginal ultrasound (Voluson™ P8; GE Healthcare, USA)-guided follicular aspiration was performed at 36 hours post-hCG administration to collect luteinizing GCs and follicular fluid from mature pre-ovulatory follicles (16-22 mm diameter). GCs were immediately processed for glycogen quantification and luteal gene expression profiling, while follicular fluid was centrifuged (3,000 ×g, 10 minutes) for subsequent progesterone quantification. The study protocol, designed in accordance with Declaration of Helsinki principles, received prior approval from the Institutional Review Board of Renmin Hospital, Wuhan University (Approval No.: WDRY2024-K262), with continuous monitoring ensuring strict adherence to ethical safeguards regarding patient autonomy and biological sample handling.

### Superovulation

Mouse superovulation was induced via intraperitoneal administration of 5 IU pregnant mare serum gonadotropin (PMSG; NSHF, China), followed by 5 IU hCG (NSHF, China) 48 hours post-PMSG administration to trigger follicle maturation and subsequent ovulation/luteinization. In sheep, estrus synchronization was achieved through intravaginal insertion of progesterone-impregnated controlled internal drug release (CIDR) devices (Zoetis, USA). Following 10 days of CIDR priming, ewes received 400 IU PMSG via intramuscular injection. CIDR removal at 12 days post-insertion coincided with 12.5 μg LH-A3 (NSHF, China) administration to induce ovulation/luteinization. Three timely subcutaneous glucose injections (1.5 g/kg body weight) were administered at 3, 12 and 18 hours post-LH-A3 treatment. Post-mating evaluation included luteal functional assessment through serum progesterone quantification 6 days post-mating and ultimate pregnancy confirmation via ultrasonography (Kxele, China) 30 days post-conception.

### Transcriptome analysis

Mouse ovaries were collected at H0, H1 and H6 for scRNA-seq (Yingzi Gene, China). Following cell viability assessment (>85%) and density calibration (1,000 cells/μL), approximately 13,000 cells *per* sample were loaded into each channel for processing on the 10 × Chromium Single Cell Platform (10 × Genomics, using the Next GEM Chip G and the Single Cell 3′ Kit v3.1), involving gel beads in emulsion (GEM) generation, reverse transcription, cDNA amplification, and Illumina-compatible library preparation. Libraries were quantified via Qubit Fluorometry (Thermo, USA) and sequenced on the Illumina Nova6000 platform (Illumina, USA) using 150-base-pair paired-end reads. Raw sequencing data underwent primary analysis using Cell Ranger (v2.1.), performing demultiplexing, genome alignment (GRCm38), Unique Molecular Identifier (UMI), and quality filtering. The first ten principal components were selected for dimensionality reduction, followed by visualization via t-Distributed Stochastic Neighbor Embedding (tSNE). To delineate cell clusters, we employed a graph-based approach that relies on the construction of a sparse nearest-neighbor network and subsequent optimization with the Louvain algorithm (3). Inter-cluster differential gene expression was assessed using the Bimod test integrated within the Seurat framework, with genes meeting a false discovery rate (FDR) ≤ 0.05 and an absolute log₂ fold change ≥ 1 considered significant. Furthermore, we analyzed spatial transcriptomic data derived from mouse ovaries. The raw count matrices were downloaded from the GEO repository under accession number GSE240271, while the curated metadata and cell annotations were sourced from the corresponding GitHub repository. (https://github.com/madhavmantri/mouse_ovulation).

While scRNA-seq enables high-resolution transcriptomic characterization of GCs, its temporal accuracy diminishes in discriminating pre-ovulatory follicle GCs across discrete post-hCG intervals. Conversely, spatial transcriptomics preserves locational fidelity but lacks the resolution of scRNA-seq. To synergize these complementary modalities, we generated a high-resolution spatiotemporal transcriptomic atlas of mouse GCs by applying established multi-omics integration methods (4, 5, 6). As part of the data preprocessing stage in our integrative workflow, we normalized the total Unique Molecular Identifier (UMI) counts from the scRNA-seq data using the *scanpy.pp.normalize_total*. Spatial cell-type mapping was achieved via a PyTorch-implemented deep neural network (*tangram.map_cells_to_space*), trained over 2,000 epochs (learning rate=0.01, Adam optimizer) on 220 ovary-specific marker genes under RNA count-based spatial constraints (*Density_prior=’rna_count_based’*). We optimized the model by minimizing the Kullback-Leibler divergence between the scRNA-seq and spatial transcriptomic datasets. This process involved projecting cellular annotations (*Subclass_label*) onto spatial coordinates via probabilistic embedding (*tg.project_cell_annotations*) and then resolving the dominant spot-level cell types using maximum likelihood estimation (*tangram_ct_pred*). Integrated transcriptomes were obtained with *tg.project_genes*, after which we implemented a multi-level quality control procedure: (i) Training convergence: visualized via loss function trajectories (*tg.plot_training_scores*); (ii) Cross-platform consistency: quantified through pairwise gene expression correlation (*tg.plot_genes_sc*); (iii) Spatial fidelity: validated via AUC metrics for marker gene colocalization (*tg.compare_spatial_geneexp*). Finally, the temporal expression dynamics of genes in GCs post-hCG administration were profiled using STEM (Short Time-series Expression Miner) with hierarchical clustering (Euclidean distance, complete linkage).

Human transcriptome data was extracted from the GEO database, with login numbers GSE133868. GO enrichment analysis of genes was performed through DAVID database (https://david.ncifcrf.gov/tools.jsp). KEGG enrichment analysis of genes were performed through KOBAS database (http://kobas.cbi.pku.edu.cn/home.d).

### Follicle culture

Small antral follicles (250-300 μm diameter) were micro dissected from mouse ovaries using 33-gauge precision needles (KONSFI, China) and subsequently cultured individually in 96-well round bottom cell culture plates (Bkmbio, 110303005, China) under standard conditions (37°C, 5% CO₂, humidified atmosphere). Follicle culture medium consisted of α-MEM (Gibco, C12571500BT, USA) supplemented with 5% FBS (Serana, FBS-AS500, Germany), 1% ITS-G (Thermo, 41400045, USA), 10 mIU/mL rFSH (GONAL-f®; sigma, USA), and 100 U/mL penicillin-streptomycin (Servicebio, G4003, China). Following 96 hours of culture, follicles reaching the pre-ovulatory stage were transitioned to high-glucose luteinization induction medium. This medium consisted of ɑ-MEM supplemented with 5mg/mL D-Glucose (MCE, HY-B0389, USA), 1.5 IU/mL rhCG (Ovidrel®; Sigma, USA), 1ng/mL Prolactin (MCE, HY-P700292, USA), 10 μM Cholesterol (MCE, HY-N0322, USA), 1% ITS-G, 5% FBS, 10 mIU/mL rFSH, 10 ng/mL hEGF (Thermo, AF-100-15, USA), and 100 U/mL penicillin/streptomycin.

### RNA interference

Lentivirus-mediated RNA interference was implemented to achieve targeted gene silencing in follicles or whole ovaries. Gene-specific small interfering RNAs (siRNAs) targeting *Ugp2* (5′-GCAAACTGAGACTGGTGGAAA-3′), *Gys1* (5′-GCCCATGTCTTCACTACCGTA-3′), and *RUNX1* (5′-CGGCAGAACTGAGAAATGCTA-3′) were designed and synthesized by Genepharma (Suzhou, China), with a commercial non-targeting siRNA (MISSION®; Sigma, USA) serving as the negative control. Next, siRNA sequences were cloned into the PLKO.1-EGFP-PURO plasmid (Genecreate, China) under the human U6 promoter. For lentiviral packaging, HEK293T cells (ATCC, USA) were co-transfected in 6-well plates with three plasmids: 0.89 μg siRNA-expressing PLKO.1-EGFP-PURO, 0.44 μg pMD2.G (VSV-G envelope; Addgene, USA), and 0.67 μg pSPAX2 (packaging plasmid; Addgene, USA) using 4 μL jetPRIME® transfection reagent (PolyPlus, 101000046, France). Viral supernatants were harvested 48 hours post-transfection, filtered through 0.45 μm Polyvinylidene Fluoride (PVDF membranes; Sigma, IPVH00010, USA), and clarified by ultracentrifugation (100,000 ×g, 2 hours; Beckman, USA).

For in vitro gene knockdown, the isolated small antral follicles were transfected with lentiviral particles (titer: 6.25 × 10^8^ particles/mL) in follicle culture medium supplemented with 10 μg/mL Polybrene (Sigma, TR-1003-G, USA) for 48 hours (37°C, 5% CO₂). Transfection was confirmed by EGFP fluorescence visualization. Transfected follicles were subsequently cultured for 48 hours to reach the pre-ovulatory stage, followed by transfer to high-glucose luteinization medium for terminal stimulation. Samples were collected at predefined intervals for analysis.

For in vivo ovarian-specific knockdown, fifteen-day-old mice were anesthetized with 1% pentobarbital sodium (50 mg/kg; Sigma, USA). A total of 5 μL high-titer lentivirus (titer: 6.25 × 10⁹ particles/mL) was bilaterally microinjected beneath the ovarian bursa using a Hamilton 701N syringe (10 μL capacity; 80330, Switzerland) coupled to a 33-gauge RN needle (Hamilton, 7803-05, Switzerland). Control cohorts received equivalent volumes of non-targeting siRNA lentivirus. Mice were allowed to recover for 5 days prior to experimental analyses to ensure stable gene knockdown.

### Metabolic flux assay

Pre-ovulatory follicles (550-600 μm diameter) were micro dissected from mouse ovaries and preconditioned in glucose-depleted Dulbecco’s Modified Eagle Medium (DMEM; Gibco, 11966-025, USA) for 2 hours to deplete endogenous glucose. Following depletion, follicles were cultured for 6 hours in luteinization medium containing 2 mg/mL uniformly labeled ¹³C₆-glucose (¹³C-glucose; MCE, HY-B0389A, USA) to achieve isotopic labeling of glycolytic intermediates. Metabolic flux analysis was performed by Biotree (Shanghai, China). GCs isolated from the follicles underwent cryogenic homogenization in liquid nitrogen, followed by suspension in 1 mL of pre-chilled (−40°C) 50% methanol. After two sequential freeze-thaw cycles (dry ice/ice bath), phase separation was achieved by adding 0.4 mL chloroform with vigorous vortexing (30 sec) and centrifugation (12,000 ×g, 15 minutes, 4°C). The polar metabolite-enriched aqueous phase was analyzed via hydrophilic interaction liquid chromatography (HILIC) on an XBridge BEH Amide column (150 × 2.1 mm, 2.5 μm; Waters, USA) coupled to a Q Exactive™ Plus Orbitrap mass spectrometer (Thermo, USA) operated in negative electrospray ionization (ESI^-^) mode. Chromatographic gradient employed: solvent A (20 mM NH₄Ac/NH₄OH in water:acetonitrile, 95:5) and solvent B (acetonitrile). Gradient program: 0-3 minutes (10% A, 90% B), 3-8 minutes (25% A, 75% B), 8-10 minutes (30% A, 70% B), 10-13 minutes (50% A, 50% B), 13-16 minutes (75% A, 25% B), 16-20.5 minutes (100% A, 0% B), 20.5-25 minutes (10% A, 90% B) at a flow rate of 0.15 mL/minutes and column temperature of 25°C. Mass spectrometric parameters included: Resolution, 140,000 (at m/z 200); Automatic gain control (AGC) target, 1 × 10⁶ ions; scan range, m/z 75–1000; Column temperature: 25°C. ¹³C-isotopologue distributions were deconvoluted using El-MAVEN v.12.0, incorporating MATLAB (v2021b)-based algorithms for natural isotopic abundance correction to enable quantitative determination of metabolic flux through central carbon pathways.

### Quantification of ¹³C-Glucose incorporated into glycogen

The isolated pre-ovulatory follicles were preconditioned in glucose-depleted DMEM for 2 hours. Subsequently, follicles were incubated for 6 hours in luteinization medium supplemented with 2 mg/mL ¹³C-glucose to facilitate isotopic incorporation into glycogen. Post-culture, GCs were isolated from follicles, homogenized in ultrapure water, and partitioned into paired aliquots. The experimental aliquot was subjected to acid hydrolysis using 6 M trifluoroacetic acid (TFA; 100°C, 4 hours) to depolymerize glycogen and release bound glucose monomers, while the paired control aliquot remained unhydrolyzed to quantify background glucose levels. Following hydrolysis, samples were equilibrated to ambient temperature (25 ± 1°C) and lyophilized under reduced pressure (300 mbar, 52°C) using a rotary evaporator (Heidolph, Germany). Dried residues were reconstituted in 0.5 M methanolic 1-phenyl-3-methyl-5-pyrazolone (PMP) solution, followed by the addition of an equimolar volume of aqueous ammonia (28%, pH 12.5) to initiate derivatization. The mixture was vortexed and incubated at 70°C for 90 minutes to form glucose-PMP derivatives. Post-derivatization, samples underwent three sequential purification cycles, each involving resuspension in 1 mL ultrapure water followed by rotary evaporation (300 mbar, 40°C) to remove residual ammonia. ¹³C-glucose-PMP adducts were then extracted via biphasic partitioning (chloroform/water; 1:1 v/v), with the aqueous phase collected for ultra-performance liquid chromatography coupled to quadrupole time-of-flight mass spectrometry (UPLC-QTOF-MS; Waters, USA) analysis. Chromatographic separation was performed on an ACQUITY C18 column (100 × 2.1 mm, 1.7 μm; Waters, USA) maintained at 45°C using a binary gradient consisting of solvent A (0.1% formic acid in water) and solvent B (acetonitrile) at a flow rate of 0.4 mL/minutes. Gradient program: 0-1 minutes (99% A, 1% B); 1-2 minutes (80% A, 20% B); 2-5 minutes (60% A, 40% B); 5-6 minutes (45% A, 55% B); 6-13 minutes (20% A, 80% B); 13-17 minutes (5% A, 95% B); 17-20 minutes (99% A, 1% B). Sample injection volume was 1 μL. Mass spectrometric detection was performed in positive electrospray ionization (ESI^+^) mode with a scan range of m/z 50-1200. Instrument parameters included: desolvation gas flow, 500 L/hour at 400°C; cone gas flow, 50 L/hour; source temperature, 100°C; capillary voltage, 1000 V; cone voltage, 30 V; acquisition rate, 0.3 sec with no inter-scan delay. The sample cone voltage was optimized using the instrument’s tune page. Accurate mass measurement and elemental composition determination for precursor and fragment ions were conducted using MassLynx 4.1 software (Waters, USA), with ¹³C-glucose of glycogen calculated as: experimental aliquot ¹³C-glucose - control aliquot ¹³C-glucose.

### Metabolite analysis

Metabolomic profiling was performed by Majorbio (Shanghai, China). GCs isolated from pre-ovulatory follicles at specified post-hCG timepoints (H0 and H6) were mixed with ice-cold methanol: water (4:1, v/v) extraction solvent (containing 0.02 mg/mL 2-chloro-L-phenylalanine as internal standard). Homogenization used a Wonbio-96c tissue grinder (wonbio, China) at 50 Hz for 6 minutes (-10°C), followed by 30-minutes ultrasonic extraction (5°C, 40 kHz) and centrifugation (13,000 ×g, 15 minutes, 4°C) to collect supernatants for injection. Analysis employed a Thermo Scientific™ Vanquish UHPLC coupled to Q Exactive™ HF-X system (Thermo, USA) with heated electrospray ionization (HESI) source operated in polarity switching mode. Chromatographic separation used an ACQUITY UPLC HSS T3 column (100 mm × 2.1 mm, 1.8 μm; Waters, USA) with mobile phase A (0.1% formic acid in water: acetonitrile, 2:98) and B (0.1% formic acid in acetonitrile) at 0.4 mL/minutes, column temperature 40°C, and injection volume 5 μL. The gradient program was: 0-2 minutes (95%A, 5%B); 2-14 minutes (5%A, 95%B); 14-14.1 minutes (95%A, 5%B); 14.1-17 minutes (95%A, 5%B). Mass spectrometry parameters: full MS resolution 120,000 (@ m/z 200), dd-MS² resolution 30,000 (@ m/z 200), scan range m/z 70-1050, spray voltage +3.5 kV/-2.8 kV, capillary temperature 400°C, sheath gas 40 arb, aux gas 10 arb, S-lens RF 55%. Quality control employed pooled QC samples injected every 10 experimental samples with retention time deviation < 0.1 minutes and peak intensity RSD < 15%. Raw data processing used Progenesis QI v3.0 (Waters, USA) for peak extraction (S/N threshold > 3), alignment (5 ppm mass tolerance), and normalization (internal standard correction). Statistical analysis on Majorbio Cloud Platform (https://cloud.majorbio.com), in GPI-anchor biosynthesis pathway annotated using KEGG database (https://www.kegg.jp/).

### Analysis of glucose uptake capacity

The glucose uptake capacity of GCs was quantified using the fluorescent D-glucose analog 2-NBDG (MCE, HY-116215, USA). Briefly, GCs were isolated from pre-ovulatory follicles at specified post-hCG timepoints, washed twice with phosphate-buffered saline (PBS; pH 7.4), and resuspended in pre-warmed (37°C) 2-NBDG working solution (100 μM in glucose-depleted DMEM). Following 30-minutes incubation at 37°C under 5% CO₂, GCs were pelleted by centrifugation (600 ×g, 5 minutes), washed twice with ice-cold PBS to terminate uptake, and finally resuspended in PBS. Fluorescence intensity of internalized 2-NBDG, which directly correlates with cellular glucose uptake capacity, was measured using a BD FACSAria™ III flow cytometer (BD Biosciences, USA) equipped with a 488-nm laser and 530/30-nm emission filter. 20,000 viable cells *per* sample were acquired. Data analysis was performed using FlowJo v10.4.0 software (BD Biosciences, USA), with results expressed as geometric mean fluorescence intensity (MFI).

### Transmission electron microscope

Pre-ovulatory follicles isolated from mouse ovaries at designated post-hCG timepoints were processed for transmission electron microscopy analysis. Follicles were primarily fixed by immersion in 2.5% (v/v) glutaraldehyde (SPI Chem, USA) at 4°C for 12 hours. Subsequently, samples were rinsed thrice with 0.1 M PBS (pH 7.4) and post-fixed with 1% (w/v) osmium tetroxide (OsO₄; SPI Chem, USA) in PBS for 2 hours at 4°C. Thereafter, specimens were dehydrated through a graded ethanol series (30%, 50%, 70%, 90%, 100%; Sinopharm, China), transitioned to propylene oxide, and infiltrated with Spurr’s epoxy resin (SPI Chem, USA) using ascending resin:propylene oxide ratios (1:3, 1:1, 3:1 v/v). Pure resin-embedded samples were polymerized at 60°C for 48 hours. Ultrathin sections (60 nm thickness) were cut using a diamond knife on a Leica UC7 ultramicrotome (Leica, Germany), mounted on 200-mesh copper grids, and double-stained with 2% (w/v) uranyl acetate (SPI Chem, USA) for 30 minutes followed by Reynolds’ lead citrate for 5 minutes. Ultrastructural examination of GCs was performed using a HT7800 transmission electron microscope (Hitachi, Japan) operated at 80 kV. Digital images were acquired using an CCD camera (Gatan, USA) with standardized exposure settings.

### PAS Staining

Glycogen deposition in tissues was assessed using Periodic Acid-Schiff Staining Kit (Sigma, 395B-1KT, USA). According to manufacturer’s specifications, ovarian samples were fixed in a 4% paraformaldehyde solution (Servicebio, G1101, China) containing 30% sucrose for 48 hours at 4°C. Post-fixation, samples were embedded in Optimal Cutting Temperature (OCT) compound (Sakura, 4583, USA). The cultured follicles were directly embedded in OCT. 6-μm-thick cryosections were prepared using a freezing microtome (Leica, Germany) maintained at -20°C. Sections were washed in PBS (3 × 5 minutes), post-fixed with 4% paraformaldehyde for 20 minutes at room temperature, and rinsed in PBS (3 × 5 minutes). Subsequently, sections were oxidized in periodic acid solution for 10 minutes, rinsed in PBS (5 minutes), stained with Schiff’s reagent for 25-30 minutes in the dark, and washed in PBS for 5 minutes. Imaging was performed using a microscope (Olympus, Japan) equipped with a digital camera under 200 × magnification.

### Immunofluorescence staining

Ovaries collected at designated post-hCG timepoints were fixed in 4% paraformaldehyde for 48 hours at 4°C, dehydrated through graded ethanol series, cleared in xylene, and embedded in paraffin. 3-μm-thick sections were prepared using a rotary microtome (Leica, Germany). Sections were deparaffinized in xylene (2 × 10 minutes), rehydrated through descending ethanol gradients, and subjected to heat-induced epitope retrieval in sodium citrate buffer (Solarbio, C1032, China) at 98°C for 30 minutes. After cooling to room temperature, sections were blocked with 10% (v/v) normal goat serum (Boster, AR0009, China) in PBS for 60 minutes at room temperature. Subsequently, sections were incubated with rabbit anti-CD34 monoclonal primary antibody (1:300 dilution; Abclonal, A19015, China) diluted in SuperKine™ Enhanced Antibody Dilution Buffer (Abbkine, BMU103, China) for 12 hours at 4°C in a humidified chamber. Following three washes in PBS (5 minutes each), sections were incubated with Alexa Fluor® 594-conjugated goat anti-rabbit IgG secondary antibody (1:200 dilution; Abclonal, AS039, China) for 60 minutes at room temperature protected from light. Nuclei were counterstained with 10 μg/mL 4′,6-diamidino-2-phenylindole (DAPI; Biosharp, BL105A, China) for 10 minutes. After final PBS washes (3 × 5 minutes), sections were mounted with antifade mountant (Servicebio, G1401, China) and imaged using a confocal laser scanning microscope (Zeiss, Germany) equipped with 405-nm (DAPI) and 561-nm (Alexa Fluor® 594) lasers. Images were acquired at 200 × magnification and processed using ZEN 2.3 lite software (Zeiss, Germany).

### Lysosome analysis

Lyso-Tracker Green (Beyotime, C1047S, China) was utilized for lysosome quantification analysis. Following the manufacturer’s protocol, GCs were isolated from pre-ovulatory follicles at specified post-hCG timepoints, washed with HBSS (Beyotime, C0218, China). Subsequently, GCs were stained with the Lyso-Tracker Green working solution and incubated at 37°C for 30 minutes. Fluorescence values were measured using a flow cytometry instrument (BD Biosciences, USA), and data analysis was conducted through FlowJo 10.4. GCs with Lyso-tracker (FITC) fluorescence > 3 × 10³, defined using a threshold slightly above the negative control maximum, were termed high Lyso-tracker fluorescence GCs.

### Analysis of lysosomal acid phosphatase activity

Lysosomal acid phosphatase activity in GCs isolated from pre-ovulatory follicles at designated post-hCG timepoints was determined using an Acid Phosphatase Activity Assay Kit (Boxbio, AKFA017C, China). According to manufacturer’s instructions, GCs were resuspended in 100 μL of lysis buffer and mechanically homogenized. Lysates were centrifuged at 8,000 ×g for 10 minutes at 4°C to obtain supernatants. Aliquots (20 μL) of the supernatant were incubated with ACP working solution for 15 minutes at 37°C. Following enzymatic reactions, ACP chromogenic substrate was added and the mixture was incubated at room temperature for 10 minutes to develop color. Absorbance at 510 nm were measured using a Spark Multimode Microplate Reader (Tecan, Switzerland). Enzymatic activities were calculated from a 6-point standard curves (0.5-8.0 μmol/mL) and expressed as U per g protein, with protein concentrations determined via bicinchoninic acid (BCA) assay kit (Servicebio, G2026, China).

### MMP assay

MMP was quantified using a JC-1-based Mitochondrial Membrane Potential Assay Kit (Solarbio, M8650, China). According to manufacturer’s instructions, isolated GCs were resuspended in pre-warmed (37°C) assay buffer and incubated with 10 μM JC-1 working solution for 20 minutes at 37°C under 5% CO₂ in the dark. Subsequently, cells were pelleted by centrifugation (600 ×g, 10 minutes), washed twice with ice-cold JC-1 staining buffer, and resuspended in fresh assay buffer. Fluorescence intensities were measured using the Spark Multimode Microplate Reader with the following settings: excitation/emission = 485/535 nm (green monomeric form) and 535/590 nm (red J-aggregates). MMP was expressed as the ratio of red (590 nm) to green (535 nm) fluorescence intensity, where decreased ratios indicate mitochondrial depolarization.

### ATP quantification

ATP levels in GCs were quantified using an ATP Determination Kit (Thermo, A22066, USA) based on firefly luciferase chemiluminescence. According to manufacturer’s instructions, isolated GCs were lysed in 60 μL ice-cold ATP extraction buffer by homogenizing followed by 5-minutes incubation on ice. Lysates were centrifuged (10,000 ×g, 4°C, 5 minutes) to remove insoluble debris. Background ATP in 96-well opaque plates (Corning, 3915, USA) was quenched by incubating with 100 μL ATP detection working solution for 5 minutes at room temperature. Subsequently, 20 μL supernatant of cell lysates was mixed with 100 μL ATP detection working solution. Chemiluminescence signals (relative light units, RLU) were immediately measured using the Spark Multimode Microplate Reader. ATP concentrations were interpolated from a 6-point standard curve (0-0.64 nmol/mL) and normalized to total protein content, with results expressed as pmol/mg protein.

### Quantification of NAD⁺/NADH ratio

The NAD⁺/NADH ratio in GCs was determined using a NAD⁺/NADH Assay Kit (Beyotime, S0175, China) based on alcohol dehydrogenase (ADH)-coupled WST-8 reduction. According to manufacturer’s instructions, GCs were lysed in 120 μL ice-cold extraction buffer and centrifuged (10,000 ×g, 4°C, 10 minutes). For total NAD (NAD⁺+NADH) quantification, 20 μL supernatant was mixed with 90 μL NADH working solution and 10 μL WST-8 chromogen, then incubated at 37°C for 30 minutes. For NADH-specific quantification, an identical aliquot was heated at 60°C for 30 minutes to decompose NAD⁺ prior to reagent addition. Post-decompose, 20 μL supernatant was mixed with 90 μL ADH working solution and 10 μL WST-8 chromogen, then incubated at 37°C for 30 minutes. Absorbance at 450 nm was measured using the Spark Multimode Microplate Reader. NAD⁺ and NADH concentrations were interpolated from an 8-point NADH standard curve (0-8 nmol/mL), with NAD⁺ calculated as: Total NAD - NADH. Results were normalized to total protein concentration.

### Quantification of glycogen, free fatty acids and triglycerides

GCs isolated from pre-ovulatory follicles at designated post-hCG timepoints were analyzed for glycogen, free fatty acids, and triglycerides using standardized biochemical assays. Glycogen content was determined with a Glycogen Assay Kit (Solarbio, BC0340, China) based on enthrone colorimetric method. According to manufacturer’s instructions, GCs were homogenized in extraction buffer, sonicated (35% amplitude, 10 sec pulse × 3; Sonics & Materials Inc, USA), and boiled for 20 minutes. After centrifugation (10,000 ×g, 4°C, 10 minutes), 60 μL supernatant was mixed with 240 μL detection buffer and incubated in boiling water for 10 minutes. The absorbance at 620 nm was measured using the Spark Multimode Microplate Reade. Free fatty acids were measured using a Free Fatty Acid Quantification Kit (Boxbio, AKFA008C, China) based on copper soap colorimetry. GCs were homogenized in lysis buffer and centrifuged (10,000 ×g, 4°C, 10 minutes). The supernatant was mixed with extraction buffer by vortexing for 10 minutes. After centrifugation (3,000 ×g, 25°C, 10 minutes), the supernatant was mixed with detection buffer and incubated in room temperature for 10 minutes. Absorbance at 550 nm was measured against palmitic acid standards (0.05-1.00 μmol/mL). Triglyceride content was assessed via Triglyceride Assay Kit (Njjcbio, A110-1-1, China) employing GPO-PAP methodology: GCs were lysed in RIPA buffer containing 1 × protease/phosphatase inhibitor (Proteintech, PR20038, China), sonicated (35% amplitude, 10 sec pulse × 3), and centrifuged (10,000 ×g, 4°C, 15 minutes). 10 μL supernatant was mixed with 200 μL enzymatic working solution and incubated at 37°C for 10 minutes. Absorbance was measured at 500 nm. All values were normalized to total protein, with results expressed as mg/g protein (Glycogen), μmol/g protein (Free Fatty Acids and triglyceride).

### Progesterone determination

Serum, follicular fluid and culture medium samples were clarified by centrifugation (3,000 ×g, 4°C, 10 minutes), while ovarian tissues were homogenized in ice-cold PBS and centrifuged (10,000 ×g, 4°C, 10 minutes). Resulting supernatants were aliquoted and stored at -80°C until analysis. Progesterone concentrations were quantified via radioimmunoassay (RIA) performed by Beijing North Institute of Biological Technology, a certified clinical testing laboratory (Beijing, China) using a commercial RIA kit with validated sensitivity (2 pg/mL). The assay employed competitive binding between endogenous progesterone and ¹²⁵I-labeled progesterone tracer for anti-progesterone polyclonal antibody binding sites. Following 1 hour incubation at 37°C, antibody-bound fractions were separated using polyethylene glycol precipitation. Radioactivity was measured for 1 minute per tube in a γ-counter (PerkinElmer, USA). Standard curves (0-100 ng/mL) were established with each run, and ovarian progesterone levels were normalized to tissue weight (ng progesterone/g ovarian tissue).

### Pharmaceutical intervention protocols

All compounds were procured from MCE. Stock solutions were prepared according to manufacturers’ specifications and diluted in appropriate vehicles immediately prior to use. In vivo inhibition of glycogenolysis: Ingliforib (HY-19396) dissolved in 10% DMSO/90% corn oil was administered via intraperitoneal injection (100 μL, 15 mg/kg) 9 hours post-hCG injection. In vitro modulation: 40 nM insulin (HY-P0035) was supplemented into the luteinization medium to activate insulin signaling, while 100 nM BMS-536924 (HY-10262) was used to suppress this signaling. MAPK pathway was activated by 10 μM C16-PAF (HY-108635), whereas suppressed by 40 μM U0126 (HY-12031A). All compounds were filter-sterilized (0.22 μm; Sigma, SLGV004SL, USA) and controls included equivalent vehicles.

### Western blotting

Total proteins were extracted using RIPA lysis buffer (Thermo, 89901, USA) supplemented with 1 × protease/phosphatase inhibitor and 1 mM PMSF (Thermo, 36978, USA) on ice. Lysates were centrifuged (10,000 ×g, 4°C, 15 minutes) and supernatants quantified via BCA assay kit. Equal protein loads (20 μg/lane) were resolved on SDS-polyacrylamide gels at 80 V (stacking) and 120 V (resolving) in Tris-Glycine-SDS buffer (25 mM Tris, 250 mM glycine, 0.1% SDS, pH 8.3) and transferred to 0.45 μm PVDF membranes at 72 V for 100 minutes in transfer buffer (48 mM Tris, 39 mM glycine, 0.037% SDS, 10% methanol). Membranes were blocked with 5% non-fat dry milk (Nestlé, Switzerland) in TBST (Servicebio, G0004, China) for 1 hour at room temperature, then incubated overnight at 4°C with primary antibodies diluted in SuperKine™ Enhanced Antibody Dilution Buffer: phospho-PKCα (CST, #9375, 1:1000), SLC2A1 (Proteintech, 21829-1-AP, 1:1000), GYS1 (CST, #3886, 1:1000), GSK3B (CST, #12456, 1:1000), phospho-GYS1 (CST, #47043, 1:1000), phospho-GSK3β (CST, #9323, 1:1000), PYGB (Proteintech, 12075-1-AP, 1:1000), UGP2 (Proteintech, 10391-1-AP, 1:800), INSR/IGF1R (Abclonal, A21984, 1:1000), phospho-INSR/IGF1R (CST, #3024, 1:800), RUNX1 (Proteintech, 25315-1-AP, 1:1000), ERK1/2 (Abclonal, A4782, 1:1000), phospho-ERK1/2 (CST, #9101, 1:1000), GAPDH (Abclonal, AC002, 1:5000). After TBST washes (3 × 10 minutes), membranes were incubated with species-matched HRP-conjugated secondary antibodies (goat anti-rabbit IgG, BF03008; goat anti-mouse IgG, BF03001; Biodragon, 1:4000) for 1 hour at 25°C. Following TBST washes (3 × 10 minutes), signals were developed with SuperSignal™ West Pico PLUS Chemiluminescence Kit (Thermo, 34580, USA) and captured using a Tanon-5200 Chemiluminescence Imager (Biotanon, China). Uncropped blots are presented in Supplementary Figure S10.

### Lectin binding assay

For lectin blotting, GCs collected at H0, H1 and H6 were homogenized in ice-cold PIERCE™ IP Lysis Buffer (Thermo, 87787, USA) supplemented with 1 × protease/phosphatase inhibitor and 1 mM PMSF. Lysates were incubated at 4°C for 30 minutes with end-over-end rotation, then centrifuged at 10,000 ×g for 15 minutes at 4°C. Supernatants were quantified using BCA assay kit. Equal protein was resolved on SDS-polyacrylamide gels at 80 V (stacking) and 120 V (resolving) in Tris-Glycine-SDS buffer and transferred to 0.45 μm PVDF membranes at 72 V for 100 minutes in transfer buffer. Membranes were blocked with 1 × Carbo-Free™ Blocking Solution (Vector, SP-5040, USA) for 1 hour at 25°C with agitation, then incubated with 1 μg/mL Biotinylated-Concanavalin A (MCE, HY-NP0174, USA)/Biotinylated-Peanut Agglutinin (MCE, HY-NP0182, USA) in blocking buffer for 1 hour at 25°C. After TBST washes, membranes were probed with Streptavidin-HRP (Thermo, N100; 1:4000 dilution in blocking buffer) for 1 hour at 25°C. Following additional TBST washes, signals were developed using SuperSignal™ West Pico PLUS Chemiluminescent Substrate and captured using a Tanon-5200 Chemiluminescence Imager with automatic gain control. Full uncropped blots are provided in Supplementary Figure S10.

For lectin flow cytometry assay, GCs were isolated from H0, H1 and H6 ovaries and washed twice with ice-cold PBS. Cells were incubated with FITC labeled MALⅠ (Vector, FL-1311, USA)–20 μg *per* 10^6^ cells concentration in 1 × Carbo Free blocking buffer for 20 minutes at room temperature. Finally, cells were diluted to 10^6^/mL using 1 × Carbo Free blocking buffer and analyzed using a Cytek® Aurora Full Spectrum Flow Cytometry (Cytek, USA) equipped with a 488-nm laser and 520-nm emission filter. 10,000 viable cells *per* sample were acquired. Data analysis was performed using FlowJo v10.4.0 software, with results expressed as geometric MFI.

### EMSA protocol

Mouse RUNX1 coding sequence (NCBI RefSeq: NM_001111021) was cloned into pcDNA3.1-3XFlag plasmid (Addgene, China) for eukaryotic expression. Flag-tagged RUNX1 protein was immunoprecipitated from HEK293T cell lysates using anti-Flag M2 affinity gel (Sigma, A2220, USA) with three wash cycles in TBST. Bound proteins were eluted with 0.1 M glycine (pH 2.7) and immediately neutralized with a neutralization buffer (1 M Tris, pH 8.5). JASPAR web-based platform (https://jaspar.elixir.no/) was employed to predict the direct binding sites of RUNX1 within the promoter regions of the *Insr* and *Igf1r*. Based on the predicted motifs, double-stranded DNA probes and corresponding site-directed mutant probes were designed, and subsequently synthesized by Genecreate (Wuhan, China). EMSA was conducted using a Chemiluminescent EMSA Kit (Beyotime, GS009, China). According to manufacturer’s instructions, biotinylated double-stranded DNA probes containing consensus RUNX-binding motifs were incubated with purified Flag-RUNX1 (100 ng) in binding buffer for 30 minutes at room temperature. Protein-DNA complexes were resolved on 6% non-denaturing polyacrylamide gels (29:1 acrylamide:bis) in TBE buffer (Beyotime, R0223, China) at 100 V for 60 minutes at 4°C. Electrophoretically transferred to Hybond-N⁺ membranes (Cytiva, RPN1510B, USA) at 380 mA for 45 minutes in TBE buffer, complexes were UV-crosslinked (120 mJ/cm², 60 ses), after blocking with the streptavidin-HRP working solution, the shift-band was detected using ECL chemiluminescent reagent by exposure on a Tanon-5200 Chemiluminescence Imager. The probe sequences are provided in Table S2.

### Luciferase reporter assay

The promoter regions of *Insr* (chr8: 3,171,461-3,172,061 bp) and *Igf1r* (chr7: 67,600,075-67,602,575 bp) were PCR-amplified from mouse genomic DNA and inserted into the PGL3-Basic luciferase reporter vector (Promega, USA) using the ClonExpress Ultra One Step Cloning Kit (Vazyme, C115-01, China). Site-directed mutagenesis of RUNX1-binding motifs (core sequence: 5′-TGT<u>G</u>GT-3′→5′-TGT<u>A</u>GT-3′) was performed using Mut Express II Fast Mutagenesis Kit (Vazyme, C214-01, China) with verification by sanger sequencing. HEK293T cells seeded in 24-well plates (1 × 10^5^ cells/well) were transfected after 24 hours using jetPRIME® reagent with the following plasmid ratio per well: 250 ng pGL3-*Insr*/pGL3-*Igf1r* (wild-type or mutant), 250 ng RUNX1-pcDNA3.1 or empty vector, and 2.5 ng pRL-TK Renilla*luciferase control vector (Promega, USA). After 24-hours incubation, cells were lysed in 100 μL Passive Lysis Buffer and analyzed using the Dual-Luciferase Reporter Assay System (Promega, E1910, USA). Firefly luciferase activity was measured first using the Spark Multimode Microplate Reader, followed by Renilla luciferase quantification after Stop & Glo® reagent addition. Promoter activity was expressed as Firefly/Renilla luminescence ratio normalized to empty vector control. Primers are listed in Supplementary Table S2.

### SEAP reporter assay

A codon-optimized SEAP gene sequence [44] was cloned into the pCDH-CMV-MCS-EF1-copGFP-T2A-Puro plasmid (GeneCreate, China), followed by lentiviral packaging in HEK293T cells and concentration by ultracentrifugation to obtain high-titer lentivirus. Small antral follicles were transfected with lentivirus (titer: 6.25 × 10⁹ particles/mL) to overexpress SEAP, with successful transfection confirmed by EGFP fluorescence. Upon reaching the pre-ovulatory stage, follicles were transferred to luteinization medium for 6 hours. Next, GCs were isolated by mechanical dissection and cultured in fresh follicle medium for another hour. Following centrifugation (3,000 ×g, 10 minutes, 4°C), the GC pellets were lysed in RIPA buffer for BCA assay, while the supernatants were aspirated for SEAP assay using Phospha-Light™ SEAP Reporter Assay kit (Thermo, T1015, USA). According to manufacturer’s instructions, the supernatants were diluted 1:1 in 1 × dilution buffer, heat-inactivated at 65 ± 0.5°C for 30 minutes, cooled to 25°C, then 50 μL sample mixed with 50 μL assay buffer (5 minutes, 25°C), followed by 50 μL reaction substrate (20 minutes, 25°C in dark). Chemiluminescence (relative light units, RLU) was measured using the Spark Multimode Microplate Reader. SEAP activity was normalized to total protein content.

### qRT-PCR assay

Total RNA was isolated using TRIzol™ Reagent (Takara, 9109, Japan) and quantified with a NanoDrop 2000 spectrophotometer (Thermo, USA). Reverse transcription was performed with 1 μg total RNA using Evo M-MLV RT Kit with gDNA clean (AGbio, AG11728, China) in 20 μL reactions (37°C for 15 minutes, 85°C for 5 sec). qPCR was conducted in triplicate on a CFX384/96 Touch™ Real-Time PCR System (Bio-Rad, USA) with 10 μL reaction volumes containing: 5 μL SYBR Green Premix Pro Taq HS (AGbio, AG11739, China), 4 μL cDNA template and 250 nM gene-specific primers (1 μL). Thermal cycling parameters: initial denaturation at 95°C for 15 minutes; 35 cycles of 95°C for 10 sec and 60°C for 30 sec (combined annealing/extension); followed by melt curve analysis (60°C to 95°C, increment 0.5°C/5 sec). Specificity was confirmed by single-peak melt curves of PCR products. Relative mRNA expression was calculated by the 2^-ΔΔCt^ method using *Actb* or *Gapdh* as endogenous controls. Amplified products were visualized through agarose gel electrophoresis (80V, 80 mA, 75 minutes). The primer sequences are provided in Table S2.

### ChIP-qPCR assay

ChIP-qPCR was performed using GCs isolated from pre-ovulatory follicles at specified post-hCG intervals (H0, H6). Cells were fixed in 10 mL DMEM/F-12 (Gibco, 11320033, USA) containing freshly prepared 1% formaldehyde (CST, 12606S, USA) for 10 minutes at room temperature with rotation (15 rpm). Fixation was quenched by adding 1 mL 1.5 M glycine followed by 5 minutes rotation. Cell pellets were washed thrice with ice-cold PBS in 1.5 mL centrifuge tubes (Axygen, MCT-150-C, USA), then lysed in pre-chilled cytomembrane lysis buffer (10 mM HEPES pH 7.9, 0.5% IGEPAL-CA630, 1.5 mM MgCl₂, 10 mM KCl, 1 × protease inhibitor cocktail [Abclonal, RM02916, China]) for 15 minutes at 4°C with vortex agitation (10 sec/5 minutes interval). Nuclear lysis was conducted in pre-chilled buffer (0.5% SDS, 10 mM EDTA, 50 mM Tris-HCl pH 8.1, 1 × protease inhibitor cocktail) for 15 minutes at 4°C. Chromatin was sonicated using a ultrasonic disintegrator (Bioruptor Plus; Belgium) with 30 sec ON/30 sec OFF cycles (30 cycles total) in 4°C water bath to generate 200-500 bp fragments validated by agarose electrophoresis. Sonicated lysates were centrifuged at 10,000 ×g for 2 minutes at 4°C, and supernatants diluted 1:10 in ChIP IP buffer (0.01% SDS, 1% Triton X-100, 2 mM EDTA, 50 mM Tris-HCl pH 8.0, 150 mM NaCl, 1 × protease inhibitor cocktail). For immunoprecipitation, pre-cleared chromatin was incubated with 5 μg anti-RUNX1 antibody (Proteintech, 25315-1-AP, China) or species-matched IgG control (Abcam, ab172730, UK) conjugated to protein A/G Dynabeads (Thermo, 26162, USA) with rotation overnight at 4°C. The beads were washed, eluted, and reverse cross-linked. DNA purification was performed using the QIAquick® PCR Purification Kit (QIAGEN, 28104, Germany). Subsequently, the DNA was utilized for qPCR analysis with specific primers. Amplified products were visualized through agarose gel electrophoresis (80V, 80 mA, 75 minutes). The primer sequences are detailed in Table S2.

### Embryo transfer

Donor female mice (KM strain, 8 weeks) underwent superovulation and mating with fertile males. Concurrently, recipient females (8 weeks) in estrus were identified via vaginal smears [45] and paired with vasectomized males. Vaginal plug confirmation at 08:00 the following morning designated gestational day (GD) 0.5. Donors were euthanized at GD 3.5 (96 hours post-hCG), and blastocysts were flushed from uterine horns using KSOM embryo culture medium (Sigma, MR-101, USA) maintained at 37°C under 5% CO₂. Pseudopregnant recipients were anesthetized with 1% pentobarbital sodium. Six morphologically intact blastocysts per uterine horn were surgically transferred into each uterine horn using a glass transfer pipette (Jitianbio, JT0061, China). Muscle and skin layers were closed with absorbable sutures (sh-Jinhuan, R611 6-0, China). Live birth outcomes were recorded daily from GD 19 until delivery completion.

### Statistics analysis

Statistical analyses were using GraphPad Prism 10.0 (GraphPad). Data were expressed as the mean ± SD. Two-tailed unpaired Student’s t test and one-way analysis of variance followed by Tukey’s post hoc test were used to analyze the statistical significance between two groups and among multiple groups, respectively. Chi-squared test was used in the comparison between the percentages. The statistical significance was set at P-value < 0.05. Most experiments were repeated at least thrice for reproducibility. The number of experimental replicates is indicated in the corresponding figure legend. If no number is listed, the experiment was conducted once.

## DATA AVAILABILITY

The single-cell RNA-seq data reported in this study have been deposited in the GEO database under accession number GSE297300. All data are available from the corresponding author upon reasonable request.

## FUNDING

This research was supported by the Biological Breeding-Major Projects in National Science and Technology, Grant/Award Number: 2023ZD04049; Fundamental Research Funds for the Central Universities (2662023DKPY001) and the National Natural Science Foundation of China (31701301).

## SUPPORTING INFORMATION

This article contains supporting information.

## ACKNOWLEDGEMENTS

We extend our deepest appreciation and respect to Mrs. Fu Bijun, Dr. He’s mother, for her genuine care and encouragement throughout this project.

## AUTHORS’ CONTRIBUTION

C.H. conceived, designed, performed, funded the experiments, analysis the data, and wrote the manuscript; J.L., Q.L., C.L., G.L., and X.L. anticipated in experiment design and conduction, data analysis, and manuscript preparation; X.W., Y.W., R.L., H.W., H.S., Y.Z., W.K., Z.R., Z.W., B.T., C.W., Q.W., G.H and S.Z assisted with sample collection and experiments conduction; C.H., J.L., Q.L., C.L., G.L., and X.L. revised this manuscript. H.C., G.L., and Q.X. supervised this project. All authors approved the final version.

## DECLARATION OF INTERESTS

The authors declare that they have no conflicts of interest with the contents of this article.

## Notes

### Competing Interest Statement

The authors have declared no competing interest.

### Summary of Updates

In this revised manuscript, we have implemented a new transcriptomic analysis method. Additionally, we have provided overall ovarian PAS staining results during luteinization and conducted a re-measurement of energy fluctuations associated with this process (Figure 4).

