## Supporting information for "Granulosa cell glycogen fuels the avascular corpus luteum"

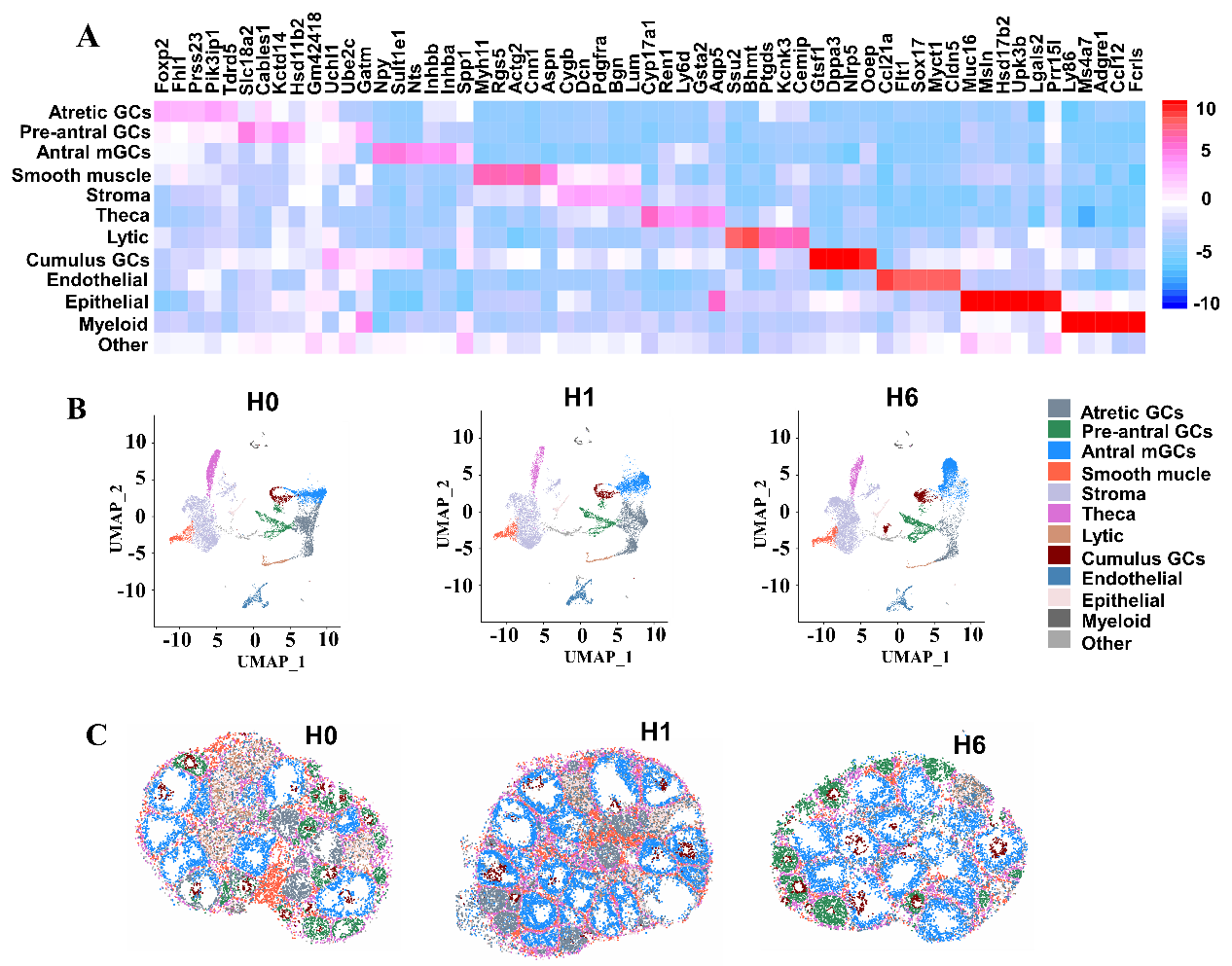

**Figure S1. Supplementary data for Figure 1**. (A) Heatmap displaying five key marker genes used for cell cluster identification. (B) UMAP visualization of single-cell RNA-seq data identifying 12 distinct cell types in H0, H1 and H6 ovaries. (C) Spatial mapping of the 12 identified cell types within ovarian tissue sections using publicly available spatial transcriptomics (H0, H1 and H6). (D) Randomly selected ovary-specific marker genes used for training demonstrate improved spatial resolution and clearer localization patterns in the integrated dataset compared to original spatial transcriptomic data.

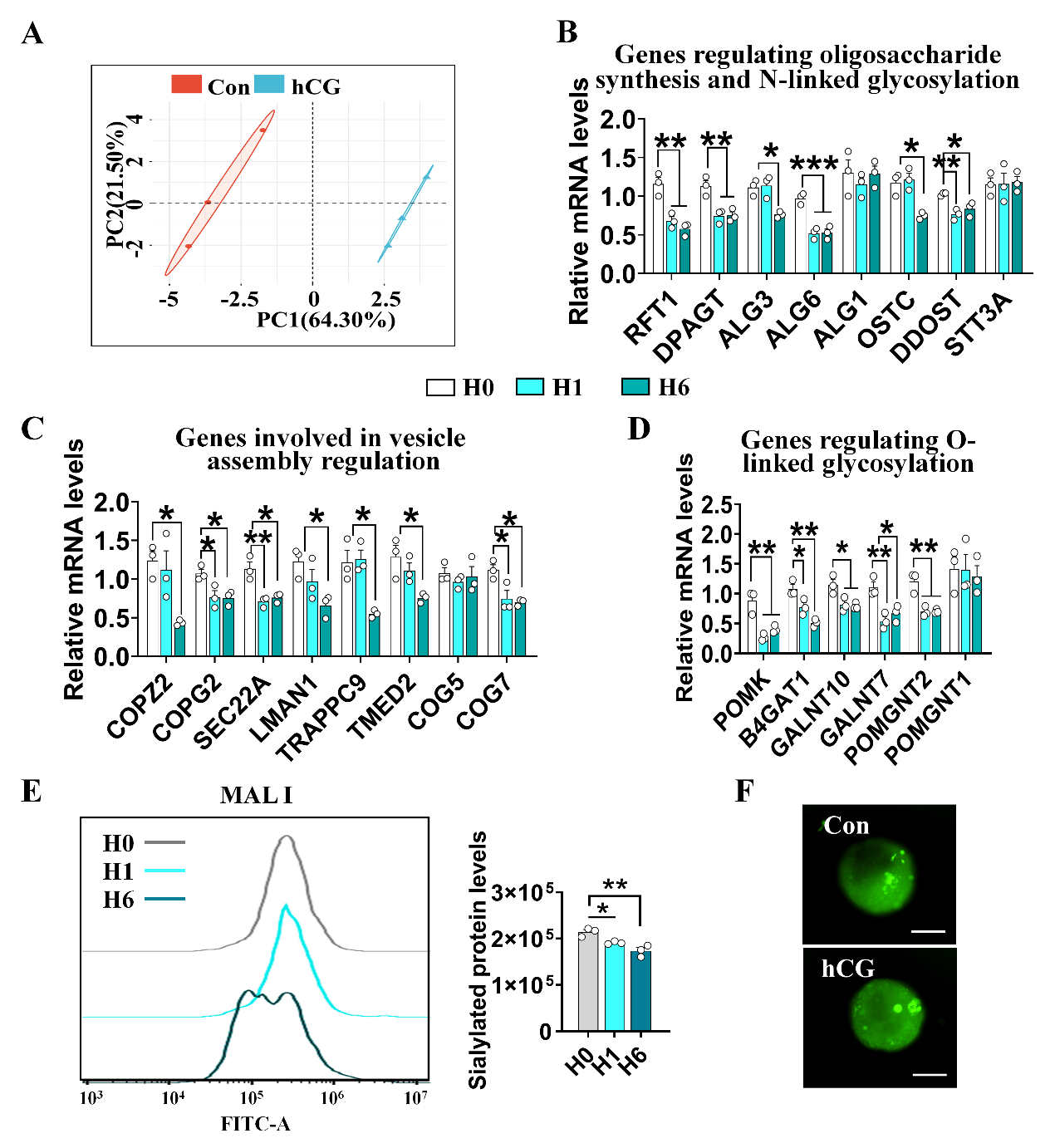

**Figure S2. Supplementary data for Figure 2.** (A) PCA of metabolic flux distributions following hCG treatment. The control was defined as the group cultured in hCG-free medium. (B-D) qRT-PCR analysis of expression changes in genes associated with oligosaccharide biosynthesis, N-linked glycosylation (B), vesicle assembly (C) and O-linked glycosylation (D) in GCs post-hCG injection (n = 3 GC samples). (E) Quantification of sialylated protein levels in GCs using maackia amurensis lectin I (MAL I) flow cytometry post-hCG injection (n = 3 GC samples). (F) Detection of green fluorescence in SEAP-expressing follicles verifies plasmid transfection efficiency. The control was defined as the group cultured in hCG-free medium. Statistical significance was assessed using one-way ANOVA followed by Tukey’s post hoc test, values are shown as mean ± SD. Significant differences are indicated by *P<0.05, **P<0.01, ***P<0.001. Results shown (B-F) are from one of at least three independent replicates; all experiments produced comparable data.

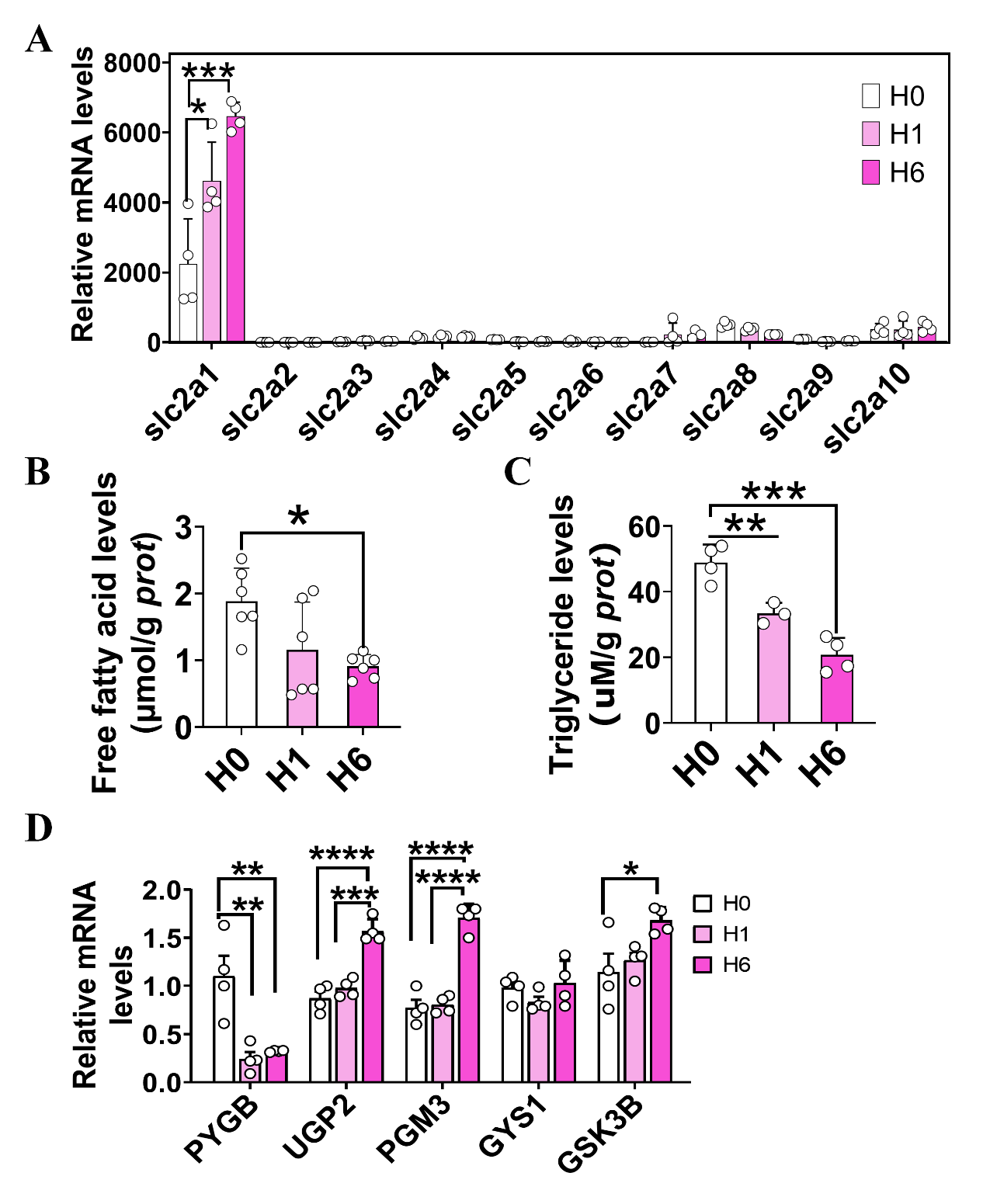

**Figure S3. Supplementary data for Figure 3.** (A) qRT-PCR analysis of glucose transporter gene expression changes post-hCG injection (n = 4 GC samples). (B) Free fatty acid content alterations post-hCG injection (n = 6 GC samples). (C) Changes in triglyceride content post-hCG injection (n = 3-4 GC samples). (D) qRT-PCR analysis of glycogen synthesis-related gene expression variations post-hCG injection (n = 4 GC samples). Statistical significance was assessed using one-way ANOVA followed by Tukey’s post hoc test, values are shown as mean ± SD. Significant differences are indicated by *P<0.05, **P<0.01, ***P<0.001, ****P<0.0001. Results shown (A-D) are from one of at least three independent replicates; all experiments produced comparable data.

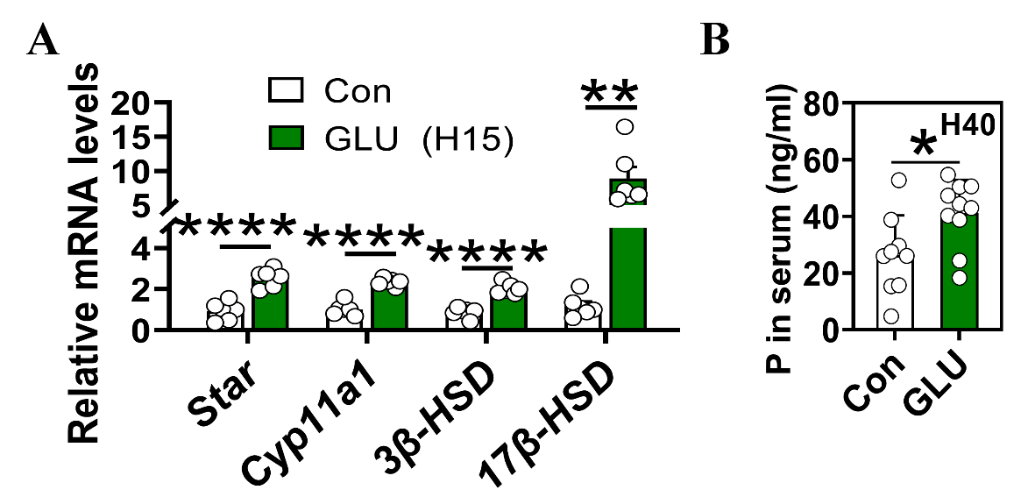

**Figure S4. Effects of glucose intake on mouse luteal function** (related to Figure 4). (A) qRT-PCR analysis of luteal gene expression changes in luteinized GCs following glucose injection at H15 (n = 6 GC samples). (B) Impact of glucose uptake on progesterone levels at H40 (n = 9-10 serum samples). Statistical significance was determined using a two-tailed unpaired Student’s t-test, values are mean ± SD. Significant differences are denoted by *P<0.05, **P<0.01, ****P<0.0001. Results shown (A, B) are from one of at least three independent replicates; all experiments produced comparable data.

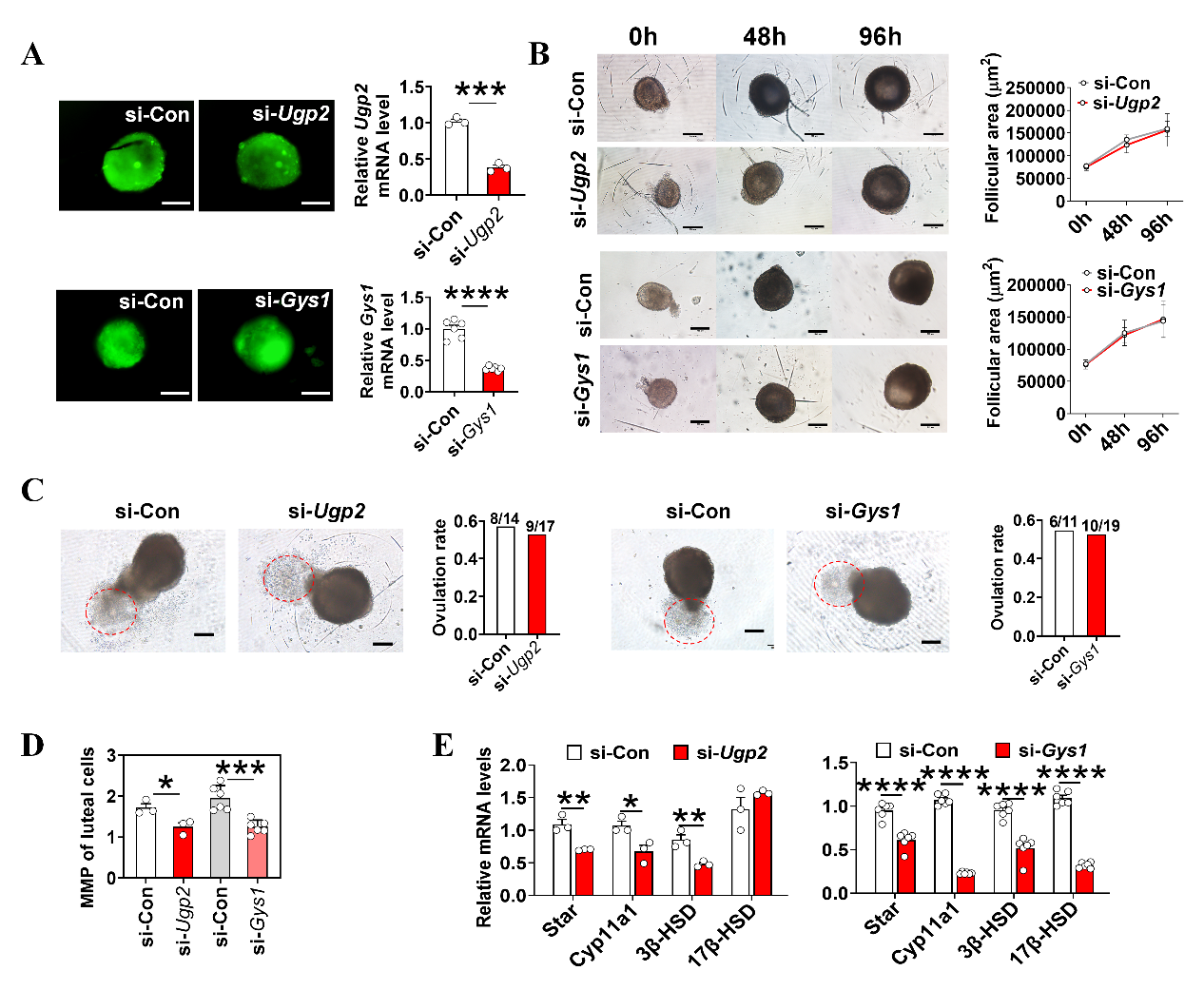

**Figure S5. Effects of *Ugp2*/*Gys1* knockdown on follicular growth, ovulation and luteinization in a follicular culture system** (related to Figure 4). (A) Verification of *Ugp2* and *Gys1* knockdown efficiency in follicular GCs, with green fluorescence confirming successful plasmid expression. (n = 3-6 GC samples), scale bar = 100 µm. (B) Effect of *Ugp2* and *Gys1* knockdown on the development of cultured follicles (n = 5-6 follicles), scale bar = 200 µm. (C) Effect of *Ugp2* and *Gys1* knockdown on the ovulatory rate of cultured follicles (n = 11-19 follicles), scale bar = 100 µm. (D) Effect of *Ugp2* and *Gys1* knockdown on mitochondrial membrane potential in luteinized GCs from cultured H15 follicles. (n = 3-6 GC samples). (E) Effect of *Ugp2* and *Gys1* knockdown on luteal gene expression in luteinized GCs from cultured H15 follicles. (n = 3-6 GC samples). Statistical significance was determined using a Chi-squared test (C, ovulatory rate) or two-tailed unpaired Student’s t-test, values are mean ± SD. Significant differences are denoted by *P<0.05, **P<0.01, ***P<0.001, ****P<0.0001. Results shown (A-E) are from one of at least three independent replicates; all experiments produced comparable data.

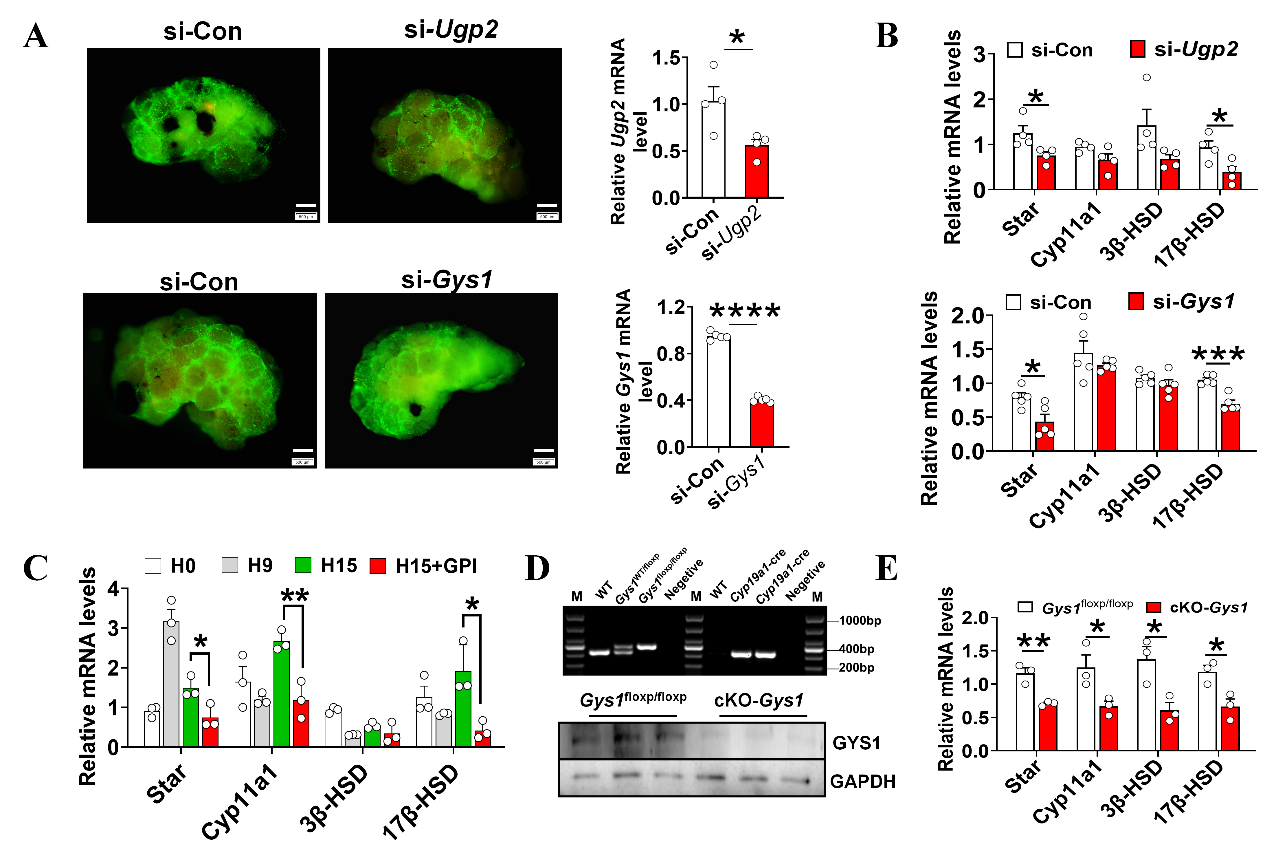

**Figure S6. Effects of *Ugp2*/*Gys1* knockdown *in vivo* on luteal function** (related to Figure 4). (A) Verification of *Ugp2* and *Gys1* knockdown efficiency *in vivo* follicles, with green fluorescence confirming successful plasmid expression (n = 4-5 GC samples), scale bar = 500 µm. (B) Alterations in luteal gene expression in H15 luteinized GCs with *Ugp2* and *Gys1* knockdown *in vivo* (n = 4-5 GC samples). (C) Effect of inhibiting glycogenolysis on luteal gene expression in H15 luteinized GCs (n = 3 GC samples). (D) PCR (Top) and Western blotting (Down) analyses confirming *Gys1* specific deletion in follicular GCs (n = 3 GC samples). (E) Effect of *Gys1* knockout on luteal gene expression in H15 luteinized GCs (n = 3 GC samples). Statistical significance was determined using a two-tailed unpaired Student’s t-test, values are mean ± SD. Significant differences are denoted by *P<0.05, **P<0.01, ***P<0.001, ****P<0.0001. Results shown (A-C) are from one of at least three independent replicates, (D, E) are from one of at least twice independent replicates; all experiments produced comparable data.

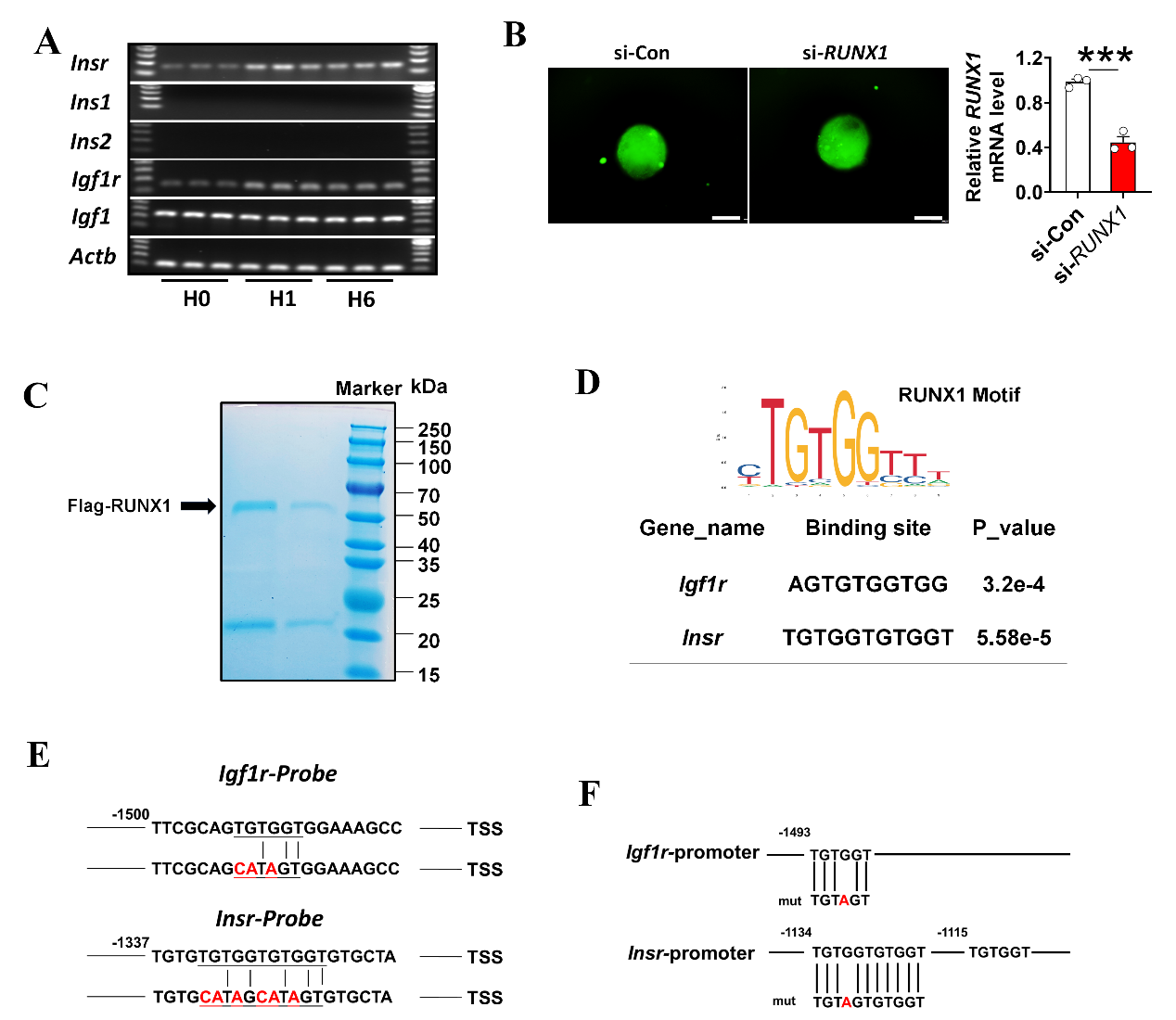

**Figure S7. Supplementary data for Figure 5.** (A) Gel electrophoresis of insulin signal-related gene expression changes post-hCG injection (n = 3 GC samples). (B) qRT-PCR validation of *RUNX1* knockdown efficiency in follicular GCs, Green fluorescence indicates successful transcription of interfering plasmids in follicles (n = 3 GC samples), scale bar = 200 µm. (C) Coomassie blue stain of Flag-RUNX1 recombinant protein. (D) Analysis based on JASPAR database showing the presence of a RUNX1 binding motif in the promoter regions of *Insr*/*Igf1r*. (E) DNA Probes of *Insr* and *Igf1r* used for the DNA-EMSA assay. (F) Mutation sites of *Insr*/*Igf1r* promoter regions for luciferase assay. Statistical significance was determined using two-tailed unpaired Student’s t-test, values are mean ± SD. Significant differences were denoted by ***P<0.001. Results shown (A, F) are from one of at least three independent replicates; all experiments produced comparable data.

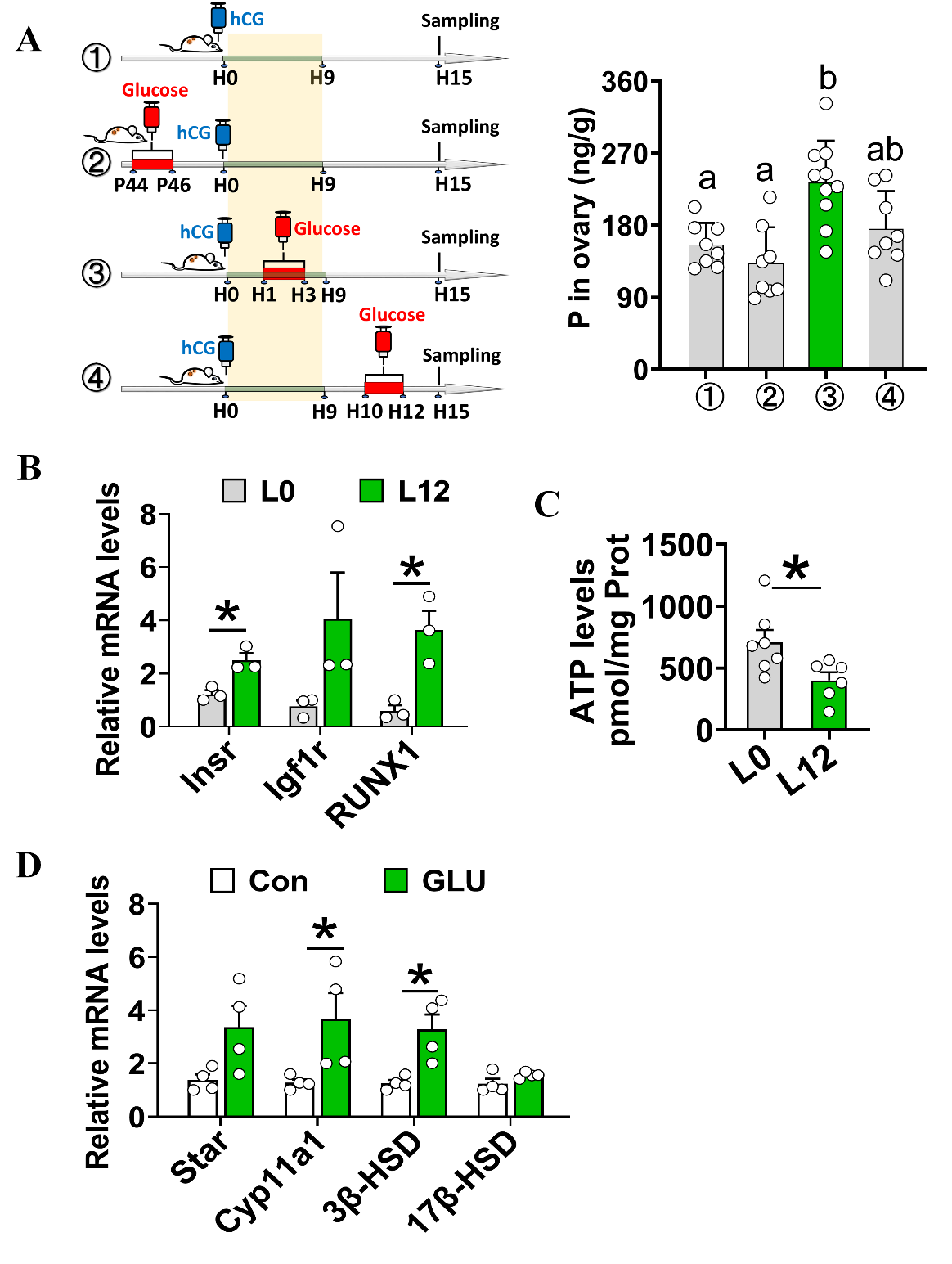

**Figure S8. Supplementary data for Figure 6.** (A) Progesterone concentrations in ovarian homogenates were quantified by radioimmunoassay following different temporal window glucose administration. Left: workflow; Right: quantification (n = 8-10 ovarian samples). (B) qRT-PCR analysis of *Insr*, *Igf1r* and *RUNX1* gene expression changes in ovine GCs post-LH injection (n = 3 GC samples). (C) Quantification of ATP content variations in ovine GCs via luciferase bioluminescence assay post-LH injection (n = 6-7 GC samples). (D) qRT-PCR analysis of luteal gene expression changes in ovine GCs post-glucose injection at GD20 (n = 4 GC samples). Statistical significance was determined using one way ANOVA with Tukey’s post hoc test (A) or two-tailed unpaired Student’s t-test, values are mean ± SD. Significant differences were denoted by *P<0.05. Results shown (A-D) are from one of at least twice independent replicates; all experiments produced comparable data.

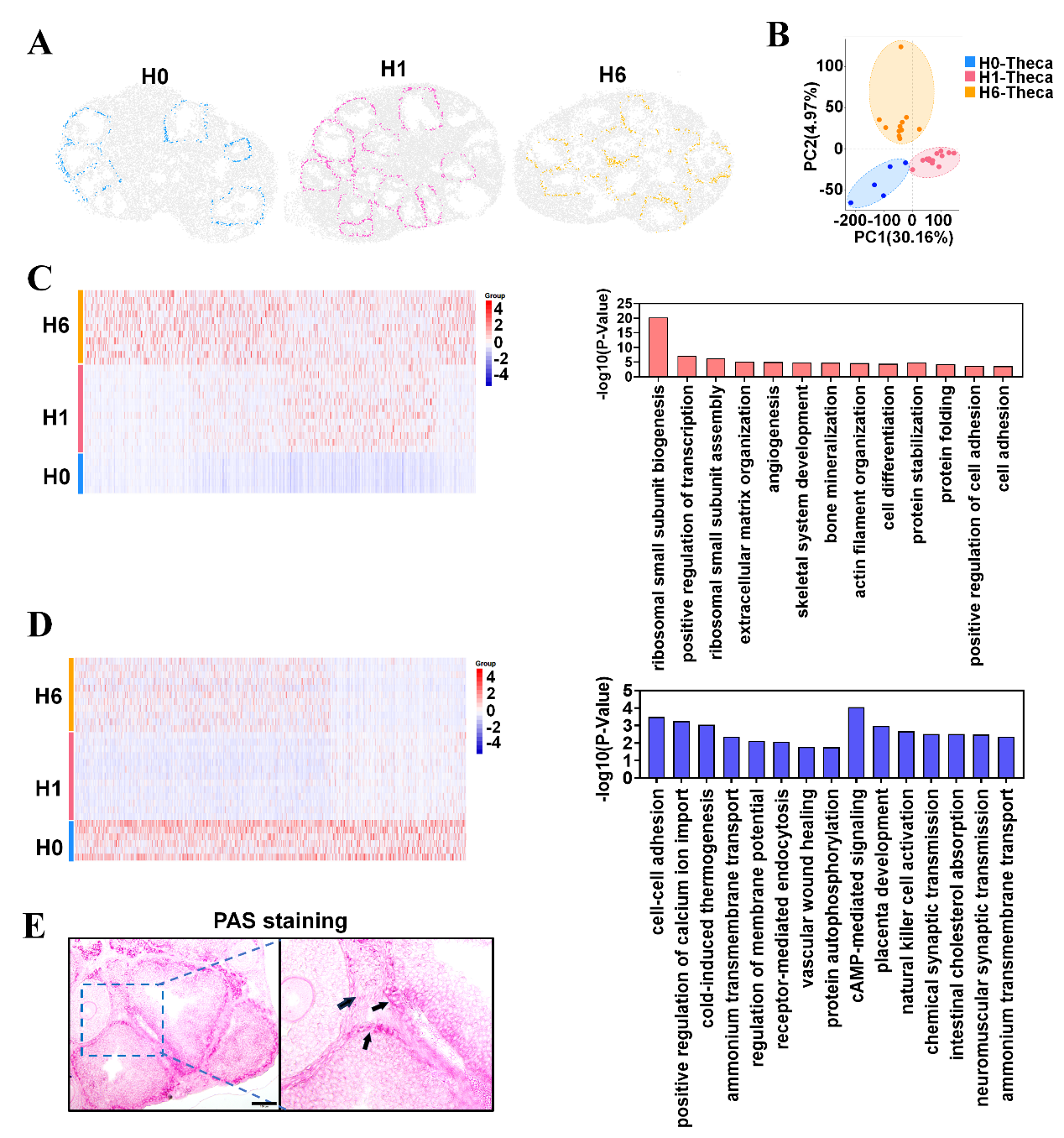

**Figure S9. Integrated multi-omics analysis of cellular response of theca cells post-hCG injection** (related to Discussion)**.** (A) Identification of theca cells in pre-ovulatory follicles at specified time points post-hCG administration using integrated analysis. (B) PCA analysis of transcriptomic differences in theca cells from pre-ovulatory follicles. (C) GO enrichment analysis of the upregulated expression cluster. Left: Heatmap of cluster-specific genes; Right: Enriched GO terms. (D) GO enrichment analysis of the downregulated expression cluster. Left: Heatmap of cluster-specific genes; Right: Enriched GO terms. (E) PAS staining of glycogen in theca cells. Scale bar = 100 μm.

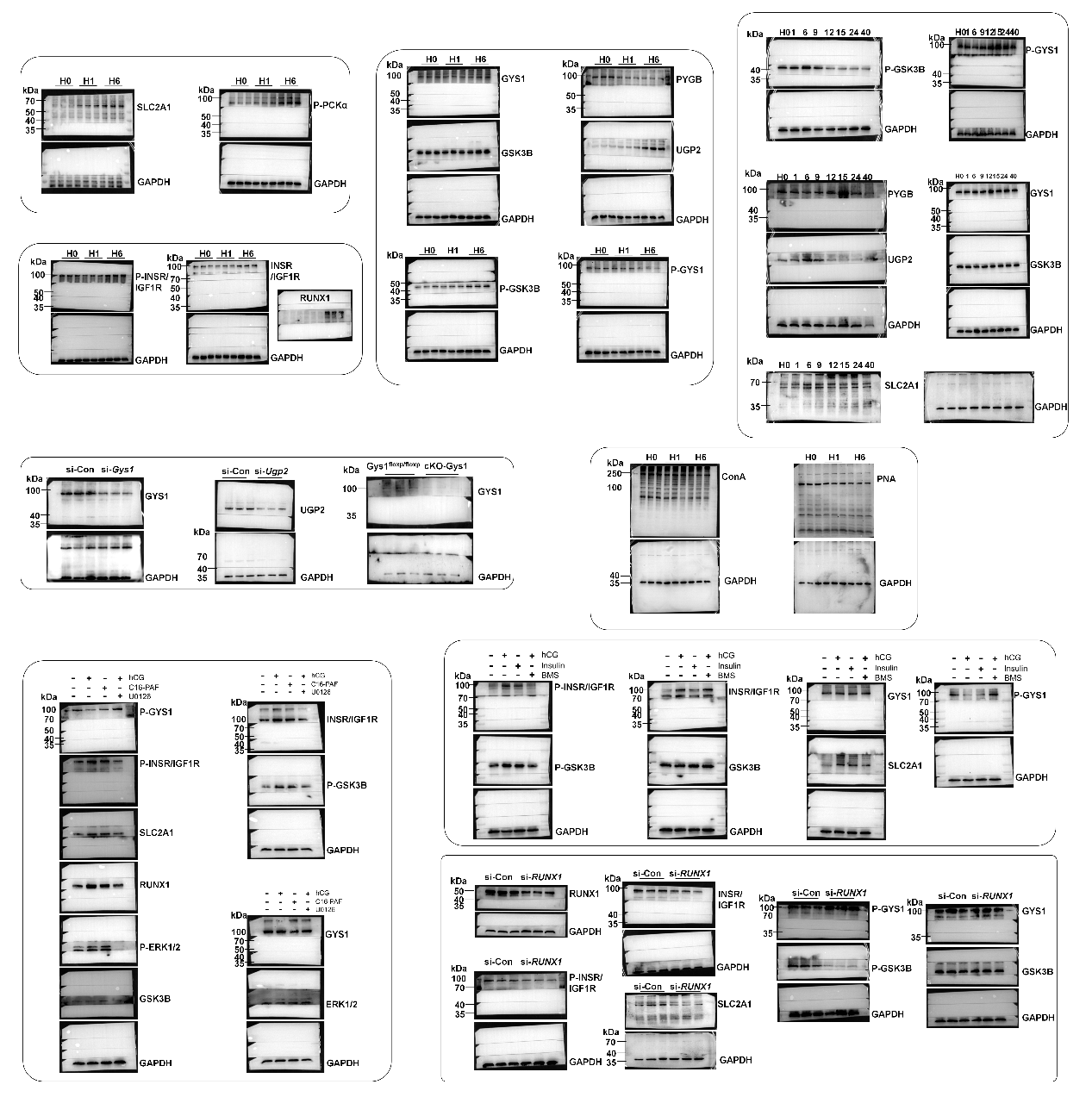

**Figure S10. Full western or lectin blots**.

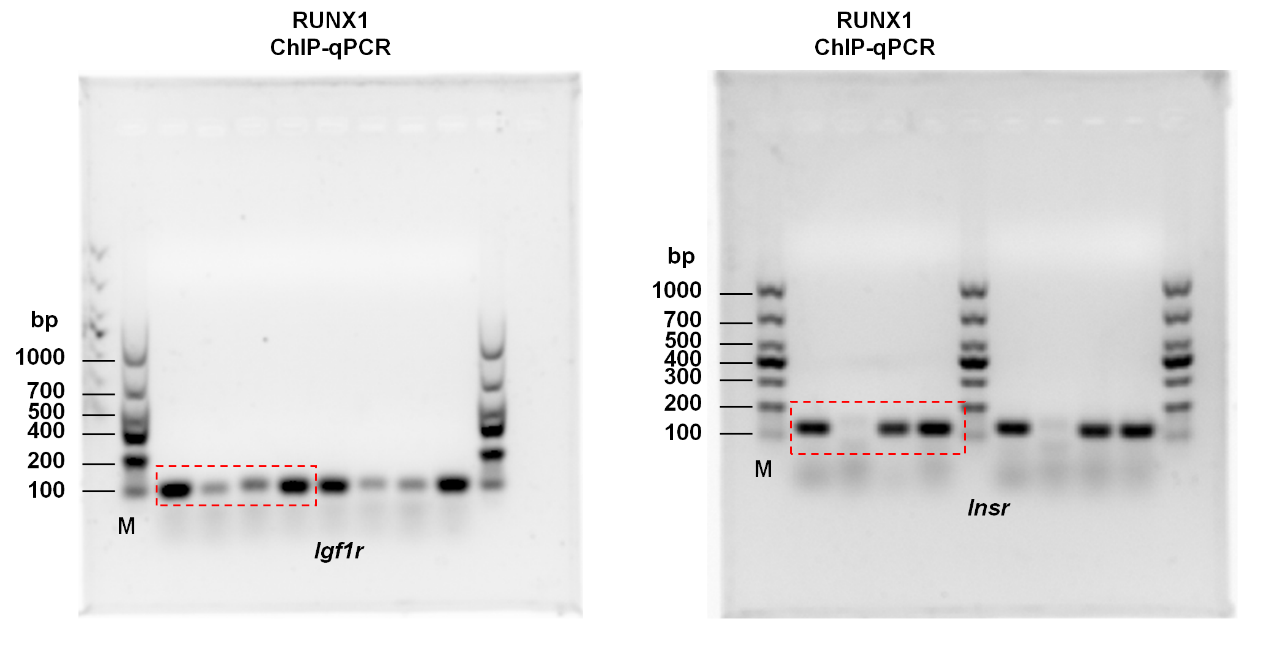

**Figure S11. Full agarose gel electrophoresis of ChIP-qPCR**

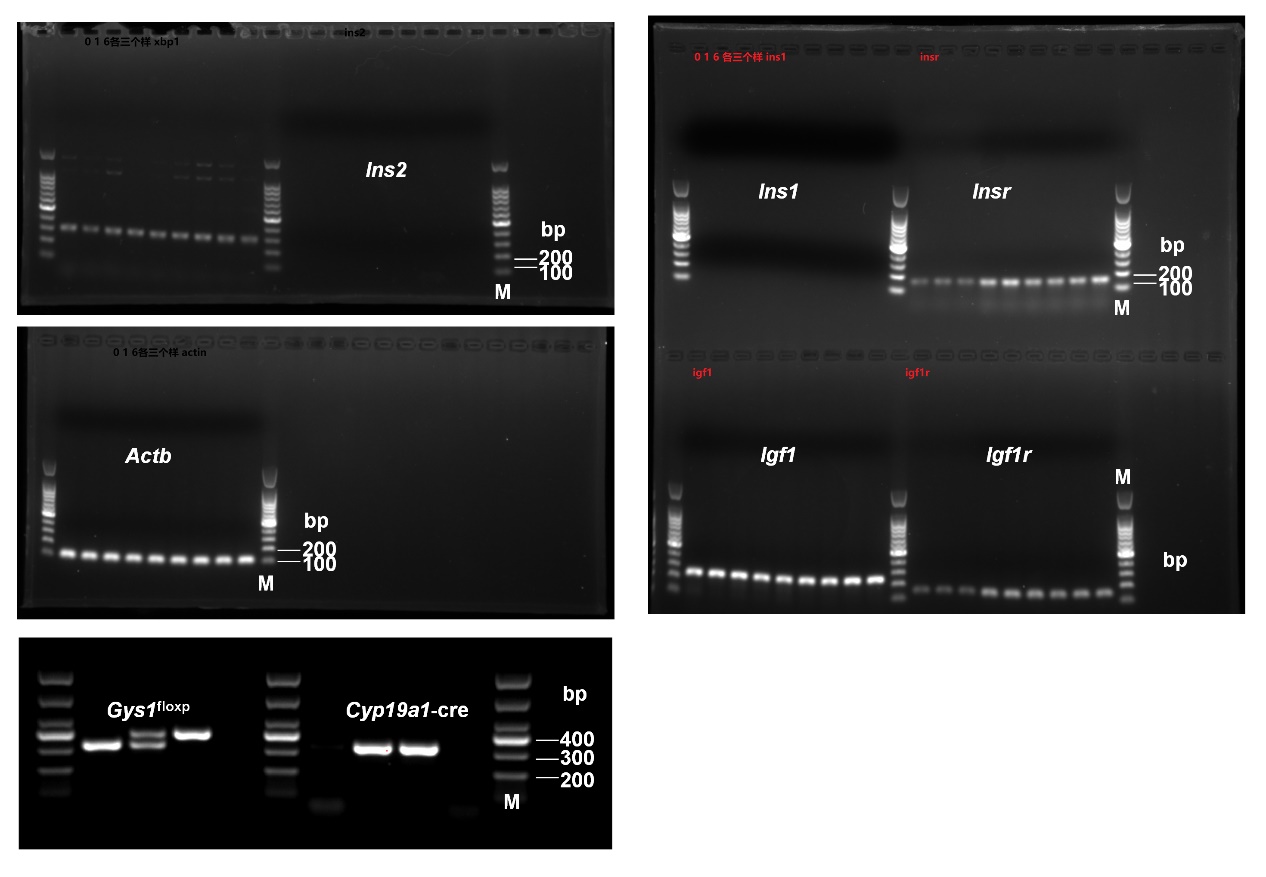

**Figure S12. Full agarose gel electrophoresis.**

**Table S1. The primers used for qPCR, ChIP-qPCR and Luciferase reporter**

| **Primer name** |  | **Primer sequence (5'-3')** |
| --- | --- | --- |
| ***Actb*** | Forward: | CCAGCCTTCCTTCTTGGGTAT |
|  | Reverse: | AGGTCTTTACGGATGTCAACG |
| ***Gapdh*** | Forward: | CGGCCGCATCTTCTTGTG |
|  | Reverse: | GTGACCAGGCGCCCAATAC |
| ***RFT1*** | Forward: | GTGGGCATAGTGAATGTCAGAC |
|  | Reverse: | AGTTAGCCACAGGAGGTTGAG |
| ***DPAGT*** | Forward: | GTGGCCCTAATAGGTGCCC |
|  | Reverse: | GCCAGCGGAGATTGAGGAC |
| ***ALG3*** | Forward: | GAACCGCGCTACACATTGC |
|  | Reverse: | GGCCTTCCAGTCAATCTCTGT |
| ***ALG6*** | Forward: | CGTGGTCTTACTAGGACTTACGG |
|  | Reverse: | GGGTAATCCAATCCCCAGTACAG |
| ***ALG1*** | Forward: | CGCTGCTGGGGTTCTACAA |
|  | Reverse: | TGTCAACTTCACAATCCGAATCC |
| ***OSTC*** | Forward: | TAGTGCTCGAATGCCCCAAC |
|  | Reverse: | CAACACTTGGAGGTTCAACGA |
| ***DDOST*** | Forward: | ACACGCACTCGCTGTTCTTC |
|  | Reverse: | ACCGACGGGGAAAAGATGATAA |
| ***STT3A*** | Forward: | TGTCGATGGCTGCTGTGTTAT |
|  | Reverse: | CTCCAATGATTCGGCCCAAAG |
| ***COPZ2*** | Forward: | GAACAGCGGGAACCTTCTCTC |
|  | Reverse: | CGGAGGGAAATGTGTCGTCATAA |
| ***COPG2*** | Forward: | GAGGAGTCTGGTAGTGGTTCC |
|  | Reverse: | TGTTCACCCTGGTTGAGTAAGT |
| ***SEC22A*** | Forward: | GTCTGCCTCAGTCATTCGTGT |
|  | Reverse: | AATGTTATAGCGCCCCGTTTT |
| ***LMAN1*** | Forward: | TGCTGTCTTTCAGCCAATTCA |
|  | Reverse: | AGCTGTATTTGTACTCGAAACGG |
| ***TRAPPC9*** | Forward: | CACCAACGGCATAAATCCTGA |
|  | Reverse: | GGCGTTCTTGTACTTGCTGTA |
| ***TMED2*** | Forward: | TGGCAATGACAGCCGTAAAG |
|  | Reverse: | TGTGTATTCTCTCCCGGACTT |
| ***COG5*** | Forward: | TTCGGGTTCCGCAGTAGTG |
|  | Reverse: | TGCTCAGCAATTACAGCTTGATG |
| ***COG7*** | Forward: | GCTTGCTGCGGAATCTCTTCA |
|  | Reverse: | GAACGATGCTACGATCTGGGG |
| ***POMK*** | Forward: | TCAGCATTGTGGCAAACGA |
|  | Reverse: | CCCCAAGAGGAAACTGGAGAC |
| ***B4GAT1*** | Forward: | TGGTTCATAAGGGATTCAAGG |
|  | Reverse: | TCAGCATCGGTGGGGAG |
| ***GALNT10*** | Forward: | AAGGAGGCTATCAGGAGGGAC |
|  | Reverse: | AGAGAGCGATTCAGGGAGATT |
| ***GALNT7*** | Forward: | TTTGCACTGTGCCGATTATAGAT |
|  | Reverse: | ACCCGTTTCCAGAGCATACTC |
| ***POMGNT2*** | Forward: | TCTTCCCTTACGCTGTCAATC |
|  | Reverse: | TGTTTCGCCAGGCTACATACT |
| ***POMGNT1*** | Forward: | GGGCTAGGAAGAAACGGAGC |
|  | Reverse: | CCCGCTGGTTTGTCAGTTTAT |
| ***IGF2R*** | Forward: | GGGAAGCTGTTGACTCCAAAA |
|  | Reverse: | GCAGCCCATAGTGGTGTTGAA |
| ***PLIN3*** | Forward: | ATGTCTAGCAATGGTACAGATGC |
|  | Reverse: | CGTGGAACTGATAAGAGGCAGG |
| ***GNPTG*** | Forward: | AACACATTCGGGCTGAATAACC |
|  | Reverse: | CCACTAGGCTAAAGCACTTGC |
| ***GNPTAB*** | Forward: | AAGTTTGGTTAGCCCAGTGAC |
|  | Reverse: | TCACCTCCTCTTGGAATCTCC |
| ***Slc2a1*** | Forward: | ATCGTCGTTGGCATCCTTAT |
|  | Reverse: | CACTCTTGGCCCGGTTCT |
| ***Slc2a2*** | Forward: | ATCGTCGTTGGCATCCTTAT |
|  | Reverse: | CACTCTTGGCCCGGTTCT |
| ***Slc2a3*** | Forward: | ATGGGGACAACGAAGGTGAC |
|  | Reverse: | GTCTCAGGTGCATTGATGACTC |
| ***Slc2a4*** | Forward: | GTGACTGGAACACTGGTCCTA |
|  | Reverse: | CCAGCCACGTTGCATTGTAG |
| ***Slc2a5*** | Forward: | CCAATATGGGTACAACGTAGCTG |
|  | Reverse: | GCGTCAAGGTGAAGGACTCAATA |
| ***Slc2a6*** | Forward: | AACCGAGGGACTCGACTATGA |
|  | Reverse: | CAAGGCATACCCAAAGCTGAA |
| ***Slc2a7*** | Forward: | CACGCACTTTGAGCGACAC |
|  | Reverse: | CCCACTTATTGACCATCAGGC |
| ***Slc2a8*** | Forward: | GCTGTCAGGGGTCAATGCT |
|  | Reverse: | GGAGTTGCTGGGGAGGCT |
| ***Slc2a9*** | Forward: | TTGCTTTAGCTTCCCTGATGTG |
|  | Reverse: | GAGAGGTTGTACCCGTAGAGG |
| ***Slc2a10*** | Forward: | ACCCATCCATCCAGTCATCA |
|  | Reverse: | TCCAAAGCCAACCGAGAA |
| ***GSK3B*** | Forward: | TGGCAGCAAGGTAACCACAG |
|  | Reverse: | CGGTTCTTAAATCGCTTGTCCTG |
| ***GYS1*** | Forward: | GAACGCAGTGCTTTTCGAGG |
|  | Reverse: | CCAGATAGTAGTTGTCACCCCAT |
| ***PYGB*** | Forward: | CCGCGACTACTTCTTCGCTC |
|  | Reverse: | CAACCCCAACTGATAAGTGGC |
| ***UGP2*** | Forward: | AGCAAAGCTATGTCTCAAGATGG |
|  | Reverse: | GAGGCTGCTGTGGTAAGTATTT |
| ***PGM3*** | Forward: | AGCAGTGGGATGCTATTTATGTC |
|  | Reverse: | TGTCTGCGCTTTCTTGTGAGT |
| ***Star*** | Forward: | GTGAAGGCTAAGGGATAA |
|  | Reverse: | TGGAGCTGGTAAGACAAC |
| ***Cyp11a1*** | Forward: | GGGCAGTTTGGAGTCAGTTTAC |
|  | Reverse: | TTTAGGACGATTCGGTCTTTCTT |
| ***3β-HSD*** | Forward: | TGGACAAAGTATTCCGACCAGA |
|  | Reverse: | GGCACACTTGCTTGAACACAG |
| ***17β-HSD*** | Forward: | CCACCTGTGTTTGGCGTGTA |
|  | Reverse: | GAGGTTGAATTGTGGATTAGGCA |
| ***Insr*** | Forward: | ATGGGCTTCGGGAGAGGAT |
|  | Reverse: | GGATGTCCATACCAGGGCAC |
| ***Igf1r*** | Forward: | GTGGGGGCTCGTGTTTCTC |
|  | Reverse: | GATCACCGTGCAGTTTTCCA |
| ***Ins1*** | Forward: | CACTTCCTACCCCTGCTGG |
|  | Reverse: | ACCACAAAGATGCTGTTTGACA |
| ***Ins2*** | Forward: | GCTTCTTCTACACACCCATGTC |
|  | Reverse: | AGCACTGATCTACAATGCCAC |
| ***Igf1*** | Forward: | CACATCATGTCGTCTTCACACC |
|  | Reverse: | GGAAGCAACACTCATCCACAATG |
| ***RUNX1*** | Forward: | GCAGGCAACGATGAAAACTACT |
|  | Reverse: | GCAACTTGTGGCGGATTTGTA |
| ***Sheep-GYS1*** | Forward: | ATGAGCCTTGGGGCTACACAC |
|  | Reverse: | GGTCTGCGATGTGTTCCTCCA |
| ***Sheep-GSK3B*** | Forward: | GGTACTGGTTCACGTCAGCA |
|  | Reverse: | AGGTCATGTTAGGGGCGTTG |
| ***Sheep-PYGB*** | Forward: | GGGGTGTTTTCACTTGCGTC |
|  | Reverse: | CCTGCCGTCTGGGTAGAATG |
| ***Sheep-UGP2*** | Forward: | TGGCCAGTTACCTAACAAATCC |
|  | Reverse: | AGTCTGTGGAAAGCATGGCA |
| ***Sheep-PGM3*** | Forward: | CGGCAAATGAGTCCCGATGA |
|  | Reverse: | TCGGGAAGACCCCTCTCATT |
| ***Sheep-Insr*** | Forward: | CTTCCTAACAACCCCTGCGA |
|  | Reverse: | AGGCTCACAGTTACGTGCTC |
| ***Sheep-Igf1r*** | Forward: | ATCGAGGGCTACCTCCACAT |
|  | Reverse: | CACGGGTGATATTCCGCAGG |
| ***Sheep-RUNX1*** | Forward: | TCTCTCGCTTCTGAAGTCCG |
|  | Reverse: | TTGCCTTAACGAAGTGCCCA |
| ***Sheep-Star*** | Forward: | CCCTGGGCATCCTCAAAGA |
|  | Reverse: | TGACACTGGGGTTCCACTCG |
| ***Sheep-Cyp11a1*** | Forward: | CGCTTTGCCTTTGAGTCCATC |
|  | Reverse: | TGAGCAGAGGGACACTGGTA |
| ***Sheep-3β-HSD*** | Forward: | TTTACAAATACCACCCCTGCTTC |
|  | Reverse: | GGTTTTCTGCTTGGCTTCCTC |
| ***Sheep-17β-HSD*** | Forward: | GAGCATAGGCGGCTTGATG |
|  | Reverse: | GTGGCGACAGTAGCGGTAGAA |
| ***Sheep-Actb*** | Forward: | CCTGCGGCATTCACGAAACTAC |
|  | Reverse: | ACAGCACCCTGTTGGCGTAGAG |
| ***Human-Star*** | Forward: | GGGAGTGGAACCCCAATGTC |
|  | Reverse: | CCAGCTCGTGAGTAATGAATGT |
| ***Human-Cyp11a1*** | Forward: | GCAGTGTCTCGGGACTTCG |
|  | Reverse: | GGCAAAGCGGAACAGGTCA |
| ***Human-3β-HSD*** | Forward: | CCAAGCTGACTGTACTTGAAGG |
|  | Reverse: | GTGTGGATGACGACCGAGAC |
| ***Human-17β-HSD*** | Forward: | GTGCTGGTGTGTAACGCAG |
|  | Reverse: | GTCCCTACTACATTCACGTCCA |
| ***Human-Actb*** | Forward: | CATGTACGTTGCTATCCAGGC |
|  | Reverse: | CTCCTTAATGTCACGCACGAT |
| ***Igf1r-RUNX1ChIP*** | Forward: | AGTCTTCAGGGGTCCCCTTT |
|  | Reverse: | AATAAGTGCATGCATGCGGC |
| ***Insr-RUNX1ChIP*** | Forward: | AGCGCTCTCTCTCTCCTCTC |
|  | Reverse: | TTAAGGTCAACCTTTGGCCT |
| ***Igf1r-EMSA*** | Forward: | GGCTTTCCACCACACTGCGAA |
|  | Reverse: | TTCGCAGTGTGGTGGAAAGCC |
| ***Igf1r-EMSAmut*** | Forward: | GGCTTTCCACTATGCTGCGAA |
|  | Reverse: | TTCGCAGCATAGTGGAAAGCC |
| ***Insr-EMSA*** | Forward: | TGTGTGTGGTGTGGTGTGCTA |
|  | Reverse: | TAGCACACCACACCACACACA |
| ***Insr-EMSAmut*** | Forward: | TGTGCATAGCATAGTGTGCTA |
|  | Reverse: | TAGCACACTATGCTATGCACA |
| ***INFU-Igf1r*** | Forward: | gagctcttacgcgtgctagcGGCTTTCCACCACACTGCG |
|  | Reverse: | acagtaccggaatgccaagcGTCTCCAACTCAGTCCTCACTCTCG |
| ***INFU-Igf1r-mut*** | Forward: | CTTTCCcatagtCTGCGAAAAAAATGCCGC |
|  | Reverse: | TCGCAGactatgGGAAAGCCCGGACGCGAC |
| ***INFU-Insr*** | Forward: | gagctcttacgcgtgctagcAAGCTCCAATCCTGATCCCC |
|  | Reverse: | acagtaccggaatgccaagcAGGGAAGGGACTGTGCGC |
| ***INFU-Insr-mut*** | Forward: | GTGTGTGtatcaTGTGGTGTGCTAGGCCAAAGG |
|  | Reverse: | CCACAtgataCACACACATAAACACACACATACAAACA |
| ***INFU-PGL3*** | Forward: | GCTTGGCATTCCGGTACTGT |
|  | Reverse: | GCTAGCACGCGTAAGAGCTC |
| **cKO-*Gys1*** | Forward: | TCAGGATTCCTAGTGAACTCCTC |
|  | Reverse: | ATCACACCCACTTTATGCCGT |
| **Cre*-Cyp19a1*** | Forward: | TACACTTTTGAGACGATTCCAGGT |
|  | Reverse: | CTAGGAATGCTCGTCAAGAAGACAG |
